# The cyanobacterium *Anabaena* uses pleomorphism as an acclimation strategy to high light stress

**DOI:** 10.64898/2026.07.01.735790

**Authors:** Cristina Velázquez-Suárez, Manuel J. Mallén-Ponce, Miguel Ángel Rubio, Mireia Burnat, José Luis Crespo, Dennis J. Nürnberg, Rocío López-Igual, Laura Corrales-Guerrero, Ignacio Luque

## Abstract

- Phytoplankton species display characteristic morphologies that are generally assumed to confer adaptive advantages, yet the functional significance of cell shape remains poorly understood. Here, we investigated whether pleomorphism contributes to acclimation to changing light environments.
- Using the cyanobacterium *Anabaena* sp. PCC 7120 as a model system, we combined molecular genetics, microscopy, physiological measurements and biophysical analyses to determine how morphology is regulated and how it affects photosynthetic performance under different light intensities.
- We show that *Anabaena* undergoes a reversible light-dependent morphological transition from rod-shaped cells under low light to large globular cells under high light stress. This transition is controlled by the relative activities of the elongasome and class A penicillin-binding proteins and is accompanied by thylakoid reorganization. The globular morphology reduces light absorption and enables cells to maintain photosynthetic activity under photoinhibitory conditions.
- Our findings establish a mechanistic link between cell-wall remodelling, cellular optics and photosynthetic performance, revealing pleomorphism as a dynamic acclimation strategy to high light stress. More broadly, this work provides experimental support for the packaging effect and highlights morphology as an active determinant of phytoplankton fitness.

## Introduction

Conservation over evolutionary time suggests that microbial morphology has a selective value (Young, 2006; Randich & Brun, 2015; Yang *et al*., 2016). However, the advantage of being a sphere instead of a rod or a helix is poorly understood and only in a few cases the suitability of a particular morphology has been elucidated (van Teeseling *et al*., 2017). Despite such constancy through time, some microbial species exhibit plasticity (*pleomorphism*), transitioning between morphologies that often correlate to environmental conditions (Yang *et al*., 2016). Cell morphology determines how microorganisms interact with the environment, i.e. by establishing the dimensions of the cell surface, or determining the presence of structures for aggregation, cell-to-cell communication, attachment, motility or predation (Young, 2006; van Teeseling *et al*., 2017). in bacteria, morphology is dictated by the cell envelope (van Teeseling *et al*., 2017), with a key contribution of the peptidoglycan (PG) cell wall. PG is a giant mesh-like polymer composed of glycosidic chains of alternating subunits of *N-*acetyl glucosamine (GlcNAc) and *N-*acetyl muramic acid (MurNAc) crosslinked by peptide bridges (Egan *et al*., 2020; Rohs & Bernhardt, 2021). As a semi-rigid structure, the cell wall limits cell enlargement, which can only occur by addition of new PG to the preexisting wall. An important concept is that PG synthesis does not occur uniformly along the cell surface; rather, the rate and precise location of PG assembly determine how the cell enlarges and ultimately dictates cell morphology (van Teeffelen & Renner, 2018; Egan *et al*., 2020; Rohs & Bernhardt, 2021).

PG assembly requires two major catalytic activities: glycosyl transferase (GT), which adds disaccharide-peptide precursors to a growing PG chain, and transpeptidase (TP), which crosslinks the pentapeptides of a nascent PG chain to those of neighboring chains. In model rod-shaped bacteria like *Escherichia coli* or *Bacillus subtilis*, PG assembly is mediated by class A penicillin-binding proteins (aPBPs)—bifunctional enzymes with GT and TP activities—and by multiprotein complexes known as the divisome and the elongasome (Egan *et al*., 2020; Rohs & Bernhardt, 2021). Among other components, these complexes contain a catalytic subunit with GT activity and a monofunctional class B penicillin-binding protein (bPBP) subunit with TP activity. Importantly, aPBPs, the elongasome and the divisome operate at specific locations. The divisome assembles at the mid-cell and catalyzes PG synthesis at the nascent septum during cell division (Du & Lutkenhaus, 2017). The elongasome directs PG synthesis at the sidewalls (i.e. the cylindrical part of the cell) allowing the cell to become longer (van Teeffelen & Renner, 2018). In *E. coli* and *B. subtilis*, the elongasome localizes in patches or bands at the sidewalls and travels short distances in the direction perpendicular to the major axis of the cell (Domínguez-Escobar *et al*., 2011; Garner *et al*., 2011; Hussain *et al*., 2018). (Shi *et al*., 2018)(Wagstaff & Löwe, 2018). aPBPs account for a substantial portion of the PG synthesis *in E. coli* (Cho *et al*., 2016; Straume *et al*., 2021), but many aspects regarding their localization and function are yet to be elucidated. Some data indicate that aPBPs can act autonomously (Cho *et al*., 2016; Lee *et al*., 2016), but other evidence support that they could act in concert with the elongasome or the divisome (Navarro *et al*., 2022). The emerging picture is that aPBPs mediate distinct modes of PG assembly at different stages of the cell cycle and at different subcellular localizations (Pazos & Vollmer, 2021). Most bacteria possess several aPBPs seemingly redundant; however, some data indicate that individual aPBPs may fulfill specialized physiological roles as shown in *E. coli* (Pazos & Vollmer, 2021).

Cyanobacteria are diderm bacteria within the Terrabacteria group (Coleman *et al*., 2021). Morphological diversity within this phylum is extensive and includes species with rod, coccoid, spiral or disk-shape morphology and many multicellular forms (Schirrmeister *et al*., 2011). Unlike other diderm bacteria, cyanobacteria contain a relatively thick PG sacculus with multiple layers (Hoiczyk & Hansel, 2000). PG synthesis complexes have been characterized to some extent in few model species (Marbouty *et al*., 2009; Xing *et al*., 2021; Velázquez-Suárez *et al*., 2022) and recent studies are uncovering novel factors and features specific for some or all species within this phylum (Corrales-Guerrero *et al*., 2018; Springstein *et al*., 2020; Xing *et al*., 2021; Velázquez-Suárez *et al*., 2023; Herrero, 2025; Ran *et al*., 2025). Multicellular cyanobacteria of the order Nostocales form multiseriate filaments of tens to hundreds of cells where the outer membrane encloses the whole filament as an external sheath, while each cell is encircled by an individual PG sacculus (Wilk *et al*., 2011). The particular complexity of their cell wall makes them interesting models for studying PG synthesis. For instance, the assembly of septal PG in these species needs to accommodate the proteinaceous channels (septal junctions) that communicate adjacent cells (Kieninger & Maldener, 2021). Further complexity arises from the ability of some species to undergo cell differentiation. This includes for example the formation of heterocysts under nitrogen-limiting conditions, which contain a special cell wall to facilitate nitrogen fixation (Zeng & Zhang, 2022).

Many features of the cyanobacterial cell structure are associated with their particular metabolism based on oxygenic photosynthesis. For instance, most species possess internal membranes, known as thylakoids, that harbor the photosynthetic apparatus (DeRuyter & Fromme, 2008). Given the central role of photosynthesis, a plethora of acclimation systems operate to tune its functioning to fluctuations in environmental factors, particularly light intensity (Wiltbank & Kehoe, 2019; Long *et al*., 2026). Excess light intensity conditions occur commonly in nature and can be particularly detrimental due to the generation of reactive oxygen species (ROS) and the associated cell damage (Tiwari *et al*., 2025). The numerous mechanisms for high light (HL) acclimation are mainly devoted to (i) limiting light absorption by reducing the antenna size or by production of photoprotective molecules (i.e carotenoids or mycosporine-like amino acids); (ii) dissipating excess excitation energy as heat, a process known as non-photochemical quenching (NPQ); (iii) redistributing excitation energy between photosystems; (iv) repairing photodamage, particularly at PSII; or (v) detoxifying ROS (Bailey & Grossman, 2008; Allahverdiyeva *et al*., 2015; Tiwari *et al*., 2025). The protection provided by the combined action of these mechanisms enables cyanobacteria to thrive under the stressful conditions imposed by HL. Previous studies have shown that cell morphology of a variety of bacterial species can be modulated by environmental factors (van Teeseling *et al*., 2017; Velázquez-Suárez *et al*., 2020). Here, we show that variations in light intensity promote reversible and dynamic changes in the morphology of the multicellular cyanobacterium *Anabaena* sp. PCC 7120 (also known as *Nostoc* sp. PCC 7120). We provide evidence that cells are rod-shaped under low light (LL) and transition into a large globular morphology upon HL exposure. This transition is achieved by post-transcriptional stimulation of aPBPs, while the elongasome is dispensable. A key feature of the large globular morphology is that it limits light absorption by pigments, probably via the so-called packaging effect. Lower absorption by large globular cells impacts photosynthesis, decreasing photoinhibition and allowing a high photosynthetic output at irradiances that are photoinhibitory for rod-shaped cells. Evidence in this article also indicates that natural *Anabaena* populations are pleomorphic, displaying a mixture of rod-shaped, large globular, and intermediate morphologies that distribute hierarchically according to incident light intensity.

## Materials and Methods

### Organisms and growth conditions

*Anabaena* sp. PCC 7120 and derived strains were routinely cultured at 30°C in BG11 medium (Rippka, 1988) in flasks illuminated with white light from Osram LED lamps 16.4K/4000K under continuous shaking or on plates of BG11 medium supplemented with 1% agar. Strains containing antibiotic resistance markers were supplemented with 10 µg ml^-1^ neomycin, 5-10 µg ml^-1^ erythromycin, 2-5 µg ml^-1^ streptomycin or 2-5 µg ml^-1^ spectinomycin as required. When indicated, cultures were grown in tubes aerated with air enriched with 1% CO2 and supplemented with 10 mM NaHCO3 as buffer.

### Plasmid and strain construction

Oligonucleotides used in this work are indicated in Table S2.

*Anabaena* mutants CSCV1 (Δ*mreB*), CSCV4 (Δ*mreC*) and CSCV2 (Δ*mreD*) described in (Velázquez-Suárez *et al*., 2020) were complemented *in trans*. In order to this, the *Anabaena* transcriptome was analyzed in search for a gene with similar expression level as the *mreBCD* operon. The *leuS* gene encoding leucyl-tRNA synthetase was selected. Constructions were made fusing 1645 bp of the 3’ region of the *leuS* gen to a linker containing an optimal Shine-Dalgarno sequence for cyanobacteria (Wei & Xia, 2019) and one of the three *mre* genes. Fusions were cloned in conjugative vectors and introduced in the corresponding Δ*mre* mutant by conjugation (Elhai & Wolk, 1988). Upon single crossover recombination at the *leuS* locus, these constructions generate an artificial operon where the *mre* gene is inserted downstream of *leuS* and expressed from the *leuS* promoter.

To generate the aPBP overexpression strains, each of the six aPBP-encoding ORF was cloned via Gibson assembly into a pDU1 based replicative vector, under the control of the strong, constitutive *trc* promoter and introduced into *Anabaena* via triparental conjugation.

The *Δall2952* mutant was constructed by deletion of a 2,244 bp internal fragment of the *all2952* ORF. DNA fragments (521 bp) flanking the sequence to be deleted were PCR amplified with primer pairs all2952-1/all2952-2 and all2952-3/all2952-4 and fused by the megaprimer PCR protocol (Burke & Barik). The resulting fragment was cloned in pSparkI (Canvax Biotech S.L., Spain), checked by Sanger sequencing and cloned as a StuI/SpeI-ended fragment into NruI/SpeI-digested pRL277 suicide vector, yielding plasmid pCSMI136, which was transferred to *Anabaena* by conjugation as described (Elhai & Wolk, 1988). Exconjugants were selected on BG11 plates containing streptomycin and spectinomycin. Selection of double recombinants was performed by culture in the presence of 5% sucrose (Cai & Wolk, 1990). Colonies were checked by PCR with primer pairs all2952-9/all2952-11 and all2952-10/all2952-11 and a clone homozygous for the inactivated *all2952* gene was selected for further work.

### Microscopy and image processing

Brightfield microscopy images were recorded with a Leica DMRE microscope with a HCX PL APO 63X NA. 1.40 oil immersion objective and a Leica Flexacam C5 camera or alternatively with an OLYMPUS BX 60 Microscope with an UPLAN FL 100X NA 1.30 oil immersion objective and a Leica DFC300 FX camera. Confocal microscopy images were obtained using an Olympus FLUOVIEW FV3000 confocal laser-scanning microscope with UPLANXAPO 40X NA 1.40 oil immersion objective or a UPLANAPO 60X NA 1.50 oil immersion HR objective. Fluorescence settings were as follows: wavelength 488 nm, detection wavelength 650 - 750 nm.

Images were processed with ImageJ/FIJI software version 2.16.0/1.54r (https://imagej.nih.gov/ij/index.html). For area and aspect ratio quantitation the following command pipeline was used: make images binary/fill holes/watershed/analyze.

### Transmission Electron Microscopy (TEM)

For cell fixation, cultures containing approximately 10 µg Chl *a* were collected, washed twice with 50 mM Tris-HCl (pH 7.5), and resuspended in 1 ml of 2.5% (v/v) glutaraldehyde in 0.1 M sodium cacodylate (pH 7.4). The solution was kept at 4°C for 2 h with occasional inversion and finally washed thrice with 0.1 M sodium cacodylate (pH 7.4). Samples were fixated in a solution of 2% KMnO4 for 2h and embedded in Spurr resin. Pictures were acquired using a Zeiss Libra 120 Transmission Electron Microscope at the University of Seville CITIUS facilities.

### RNA preparation and gene expression analyses

Total RNA isolation and northern assays were carried out as described in (Sarasa-Buisan *et al*., 2024). Probes for specific genes were generated by PCR using oligonucleotides described in Table S2.

RNA samples for RT-qPCR were extracted using the Monarch Spin RNA Isolation Kit (NEB, #T2110S) with some modifications. A culture volume equivalent to 50 µg of chlorophyll a was collected by centrifugation and lysed following the procedure previously described in the “RNA preparation” section, before proceeding with the kit according to the manufacturer’s instructions. The integrity of the extracted RNA was verified via agarose gel electrophoresis, and the absence of genomic DNA (gDNA) contamination was confirmed by PCR.

For cDNA synthesis, 0.5 µg of extracted RNA was reverse-transcribed using the iScript cDNA Synthesis Kit (BioRad, #1708890) using random hexamers as primers. The qPCR reaction was carried out using iTaq Universal SYBR Green Supermix (Bio-Rad) and 5 ng cDNA per reaction and measured in a CFX96 Touch Real-Time PCR Detection System (Bio-Rad). Probes for specific genes were generated by PCR using oligonucleotides described in Suppl. Table 1. The *rnpB* gene (Napolitano *et al*., 2012) was used as a standard gene for normalization, and expression level was expressed as 2^ΔΔCT^ between WT and over-expression strains. Oligonucleotides used as primers for qPCR are indicated in Table S2.

### Pigment analyses and absorption spectra

Chlorophyll was determined in methanolic extracts as described by Mackinney (Mackinney, 1941). Phycobiliproteins were determined as described by Siegelman and Kycia (Siegelman & Kycia, 1978). Carotenoid were extracted with acetone as described by Davies (Davies, 1976). Absorption spectra were recorded using a Thermo Scientific Genesys 180 spectrophotometer. For cell disruption, cell suspensions were supplemented with 1 mg ml^-1^ lysozyme, incubated at 30°C in a rocking platform for 1 h and sonicated for 1 minute in a Branson 250 Digital Sonifier. Full cell disruption was checked by microscopy and absorption spectra were recorded as described above.

### Oxygen evolution measurements

Oxygen evolution was measured in cell suspensions using a Clark-type oxygen electrode (Hansatech) maintained at 30 °C. Cells were collected and resuspended in BG11 medium at 4-6 μg Chl ml⁻¹. Photosynthetic activity was recorded under illumination (50 μmol photons·m⁻²·s⁻¹), while respiratory rates were measured in the dark, each for a duration of 10 minutes.

### Chlorophyll fluorometry

Chlorophyll fluorescence was measured using pulse-amplitude modulation (PAM) fluorometry with a Dual-PAM-100 (Walz) on intact cells at room temperature (Mallén-Ponce *et al*., 2021) following the recommendations by (Ogawa *et al*., 2017). Prior to measurements, cells at 4-6 μg Chl ml⁻¹ were dark-adapted for 10 minutes. To determine basal fluorescence(F₀) in in weak blue light, maximal fluorescence under actinic light (Fmʹ), and maximum fluorescence in the presence of 20 μM DCMU (FmDCMU), a 250-ms saturation pulse (5000 µmol photons m⁻² s⁻¹) was applied. The effective quantum yield of PSII [Y(II)] was calculated using the equation Y(II) = (Fmʹ – Fs)/Fmʹ, where Fs represents the steady-state fluorescence under actinic light.

### P700 measurements

Changes in P700 absorption were monitored using the Dual-PAM-100, following the procedure described in (Mallén-Ponce *et al*., 2021). The maximal photo-oxidizable P700 (Pm) was determined by applying a saturation pulse under far-red illumination (720 nm, 75 W·m⁻²).

## Results

### Light intensity dictates cell shape in Anabaena

Given the prominent role of light in cyanobacterial physiology, we tested its possible influence on *Anabaena* morphology. Throughout this work, two different light conditions were used: a dim light intensity of 20 µmol photons m^-2^ s^-1^, referred to as “low light” (LL), and a brighter light intensity of 500 µmol photons m^-2^ s^-1^, termed “high light” (HL). While cells growing under LL displayed a typical rod shape (Fig. 1A), transfer to HL conditions caused cell enlargement and transformation into a globular shape. Cell morphology was quantified by measuring two parameters, the *cell area*, which is the area encircled by the cell envelope in brightfield microscopy images, and the *aspect ratio,* calculated as the length of the axis parallel to the filament divided by the length of the axis perpendicular to the filament (Fig. 1B). The cell area showed an increase upon transfer to HL, reaching a maximum at 72 h and stabilizing thereafter. The aspect ratio was higher than 1 for cells growing in LL, corresponding to rod-shaped cells, but in HL progressively decreased to values below 1 that define cells with a globular shape (Fig. 1AB). This morphological shift was accomplished by an increase of the axis perpendicular to the filament (from an average of 2.8 to 5.3 µm) and a modest increase of the axis parallel to the filament (from an average of 4 to 4.2 µm). Intermediate morphologies between rod and globular shape were observed when cells growing in LL were transferred to light intensities ranging from 20 to 200 µmol photons m^-2^ s^-1^ (Fig 1C). Reversion into a rod-shape was observed when globular cells growing in HL were transferred back to LL, but this process was slower (Fig S1A).

**Fig. 1.**
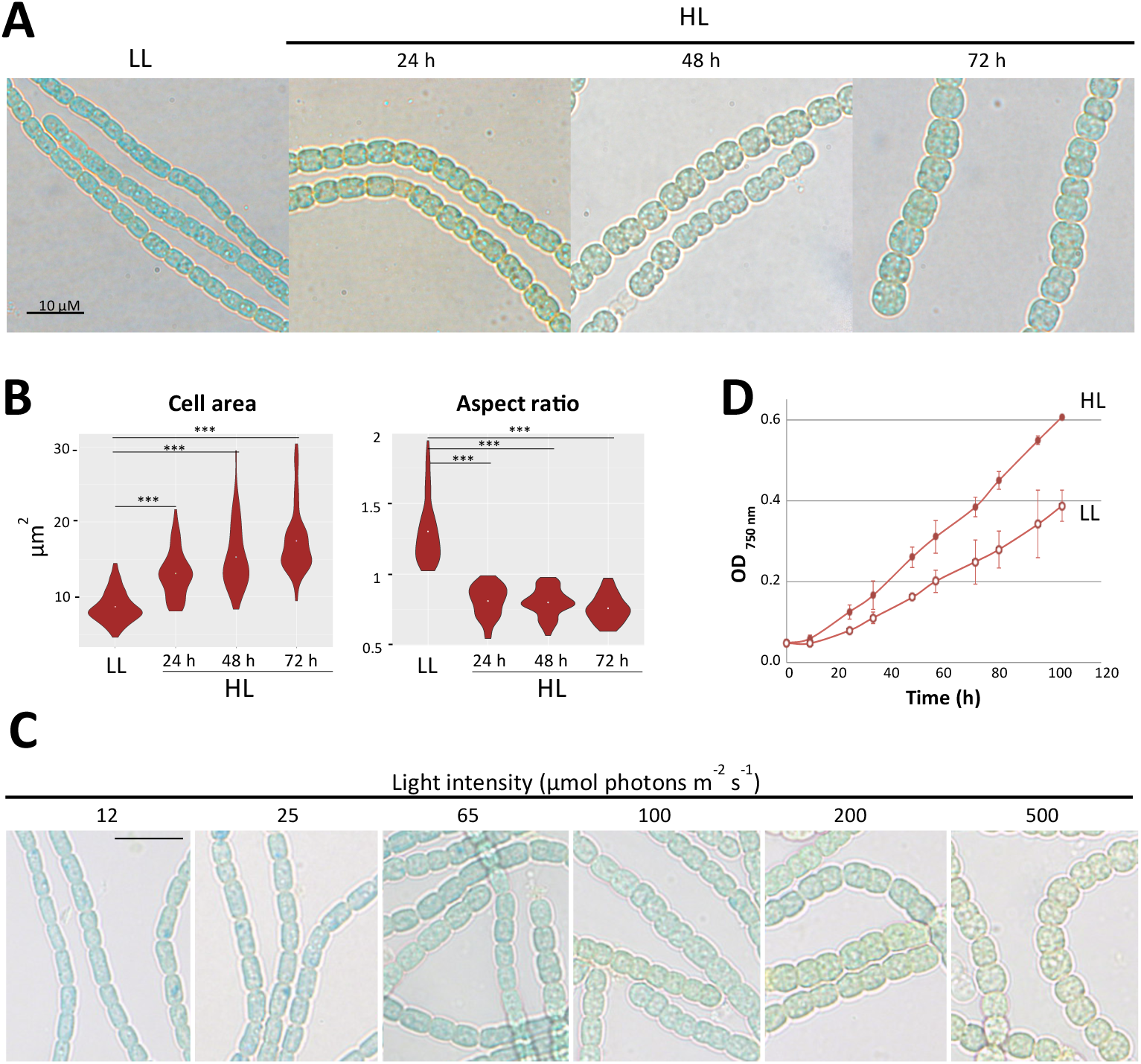
Morphological change upon transfer to HL. (A) Liquid cultures of *Anabaena* growing in LL (20 µmol photons m-2 s-1) were transferred to HL conditions (500 µmol photons m-2 s-1) for the indicated time (in hours) and observed by light microscopy. (B) Kernel density plots represent the cell area and the aspect ratio of cells cultured as in A (*n*=50 cells). The significance of differences was tested using the Welch’s-*t* test (*** means *p* < 0.001) (C) Images of *Anabaena* cells cultured in LL and transferred to the indicated light intensities for 72 h. Scale bar, 10 µm. (D) Growth curves of *Anabaena* liquid cultures incubated in LL or HL. Average values ± SD of three independent experiments are shown.

The correlation between cell size and growth rate is a well-established phenomenon in some bacterial species (Schaechter *et al*., 1958). Since *Anabaena* growth was faster in HL than in LL (Fig. 1D), we investigated whether the observed morphological shift was associated with rapid growth. To test this, LL cultures were supplemented with 1% CO2. Under these conditions, growth was significantly faster than in LL and even surpassed that of HL (Fig. S1B). However, cell morphology remained unchanged, indicating that the shift observed in HL is specific to those conditions (Fig. S1C, D).

### Role of the elongasome for the morphological transition in HL

The observed distortion of the cell sidewalls upon transfer to HL suggested a possible involvement of the elongasome. To investigate this, we took advantage of the fact that knock out mutants of several elongasome components are viable in *Anabaena* (Burnat *et al*., 2014; Velázquez-Suárez *et al*., 2020). As described, mutants of the *mreB*, *mreC* or *mreD* genes encoding major constituents of the elongasome, exhibited in LL a discoidal morphology, with cells less elongated than those of the WT (38). After 48 h of exposure to HL, cells of the three *mre* mutants were larger and adopted a globular morphology similar to the WT, although cells of the *mreD* mutant were somewhat larger (Fig. S2A, B). The low magnitude of the Cohen’s *d* values and particularly its decrease when comparing the aspect ratios of the WT and the mutants (Fig. S2C), corroborated a clear morphological convergence of all strains after 48 h in HL, despite *p* values remaining significant. Consequently, these observations suggest that the morphological shift observed in HL can occur even in the absence of the elongasome, and therefore it is not the primary driver of the PG growth in globular cells.

An unexpected observation in this experiment was that the *mreC* mutant invariably ceased growth few hours after the transfer to HL (Fig. S2D). *mreB* and *mreD* mutants were also impaired in HL, their growth showing extensive variability in different experimental replica (Fig. S2D, right plot). Genetic complementation restored both rod-shaped morphology in LL–albeit partially for the *mreD* strain–and growth in HL for all three *mre* mutants, (Fig. S3). We investigated the basis for growth impairment of these mutants in HL. Culture with CO2-enriched air ruled out that growth impairment in HL derived from an inability of the *mre* mutants to cope with a fast growth rate (Fig. S4). Of note, all three mutants kept a constant discoidal morphology in CO2-enriched cultures.

We also analyzed the functioning of photosynthesis, as acclimation is crucial for survival in HL. Results did not support that growth impairment of the *mre* mutants resulted from defective acclimation to HL (see a thorough description of these experiments in File S1). Notably, we observed that after 9 hours in HL, the *mreC* mutant accumulated 2-fold higher concentration of ROS than the wild-type, evidencing an impairment to cope with photooxidative stress, which may contribute to its reduced viability in HL. In summary, these observations reveal that despite the elongasome being dispensable for the morphological shift, its Mre subunits are required for growth in HL, MreC being particularly important under these conditions.

### Mecillinam induces Anabaena cells to become globular

Mecillinam, a β-lactam antibiotic, induce a globular morphology in a variety of rod-shaped bacterial species, including *E. coli,* prior to killing (Cho *et al*., 2014). We noticed that treatment of *Anabaena* cultures with mecillinam induced cells to become larger, adopting a globular morphology very similar to that observed in HL (Fig. 2). We reasoned that although HL and mecillinam treatment are unrelated conditions probably coupled to dissimilar responses, it was possible that they converge in some steps leading to this particular morphology. So, elucidating the events triggered by mecillinam could provide information on the mechanisms operating in HL. In rod-shaped model bacteria (*E. coli* and *B. subtilis*), mecillinam has been described to block the transpeptidase activity of the elongasome by specific inhibition of PBP2 activity (Cho *et al*., 2014). Thus, we investigated whether the morphological shift induced by mecillinam was linked to the inhibition of PBP2 in *Anabaena*. In order to this, we used the *pbp2* (*alr5045*) knock out mutant described by Burnat *et al*. (Burnat *et al*., 2014). Similar to other elongasome defective variants, cells of this mutant displayed a typical discoidal morphology in LL, with cells being less elongated than the WT (Fig. 2). Upon treatment with mecillinam, both *pbp2* mutant and wild-type cells adopted a globular morphology, becoming morphologically indistinguishable, indicating that the morphological shift was not a consequence of PBP2 inhibition (Fig. 2). In apparent consistency, we observed that cells of the three *mre* mutants also adopted a globular morphology upon treatment with mecillinam (Fig. S5). Therefore, in *Anabaena,* the transition to a globular morphology triggered by mecillinam is seemingly independent of the functional state of the elongasome.

**Fig. 2.**
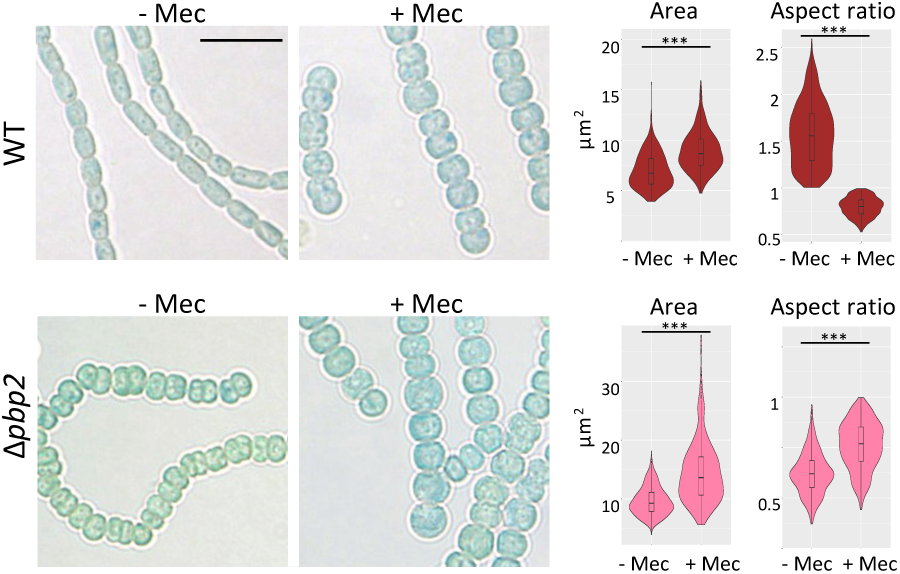
Impact of mecillinam on cell morphology. Cultures of WT *Anabaena* or the Δ*pbp2* mutant containing 1 µg Chla ml-1 were supplemented or not with 40 µg ml-1 mecillinam, as indicated, incubated for 72 hours in LL and observed by light microscopy. All pictures were taken with the same magnification (Scale bar 10 µm). Diagrams at the left side represent the cell area and the aspect ratio of cells from this experiment. The experiment was repeated three times and cells from the three experimental replica were quantitated (*n*=500). Results were statistically analized by Welch’s-*t* test (***, *p*<0.001).

### Role of class A PBPs

Given the apparent dispensability of the elongasome for the transition into a large globular form, we hypothesized the involvement of aPBPs. To investigate this, we generated strains overexpressing each of the six aPBPs encoded by the *Anabaena* genome (Leganés *et al*., 2005). Each of these strains carried a replicative plasmid with the corresponding gene downstream of a strong promoter. According to qPCR results, overexpression was successful in all cases, each gene reaching expression levels higher than those in the WT (Table S1). Of note, cells overexpressing All2952 (strain OE-*all2952*), and to a lesser extent those overexpressing Alr5326 (strain OE-*alr5326*), were larger and rounder than WT cells, while strains overexpressing other PBPs did not show morphological alterations (Fig 3A). Most importantly, cells of the OE-*all2952* strain displayed a globular morphology comparable to that of wild-type cells incubated in HL or with mecillinam (Fig. 3B, C).

**Fig. 3.**
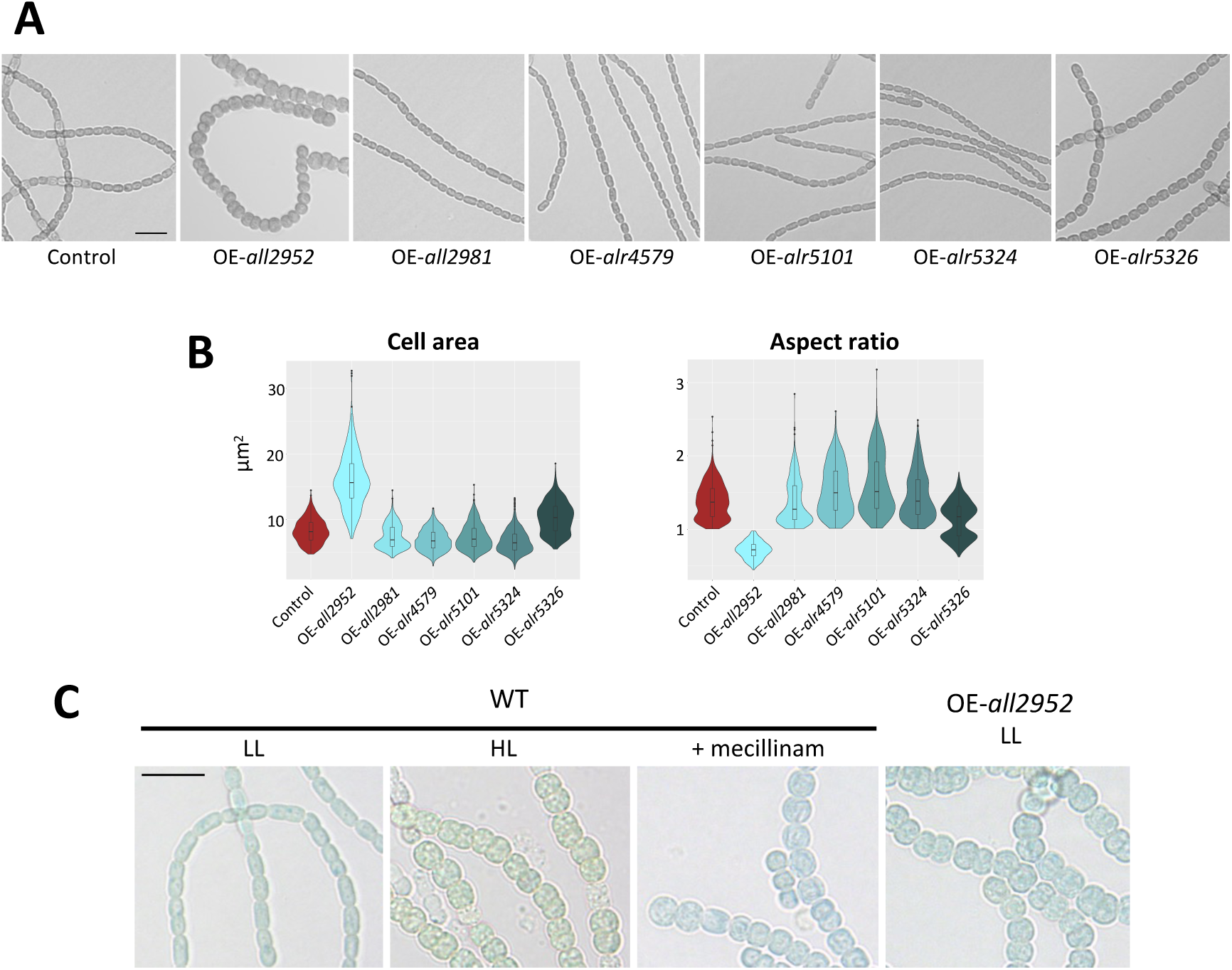
Overexpression of genes encoding aPBPs. (A) Confocal brightfield images of WT *Anabaena* and the indicated strains. Scale bar, 10 µm. (B) Diagrams represent the cell area and the aspect ratio of the indicated strains (*n*=500 cells). (C) Brightfield images of the *Anabaena* WT strain and the OE-*all2952* strain under the indicated conditions. All pictures were taken at the same magnification for comparison. Scale bar, 10 µm.

This suggested that the overabundance of some aPBPs was sufficient to induce a globular morphology, which prompted us to check whether transcription of genes encoding aPBPs was induced in HL. Northern and qPCR assays showed that in LL transcript levels of these genes were quite dissimilar. Nonetheless, as shown in Fig. S6 none of their transcripts increased in HL, ruling out transcriptional regulation. Hence, we postulated that HL would influence aPBPs through a post-transcriptional mechanism (i.e. by stimulation of aPBP activity). To test this, the OE-*all2952* strain was challenged with HL and was observed to form giant globular cells that eventually lysed, indicating massive peripheral PG synthesis, which would be consistent with HL-mediated stimulation of the large aPBP load in this strain (Fig. 4A, Fig. S7A). Interestingly, OE-*all2952* cells treated with mecillinam also became giant, further supporting that mecillinam and HL share common mechanisms regulating cell morphology (Fig. 4B, Fig. S7B).

**Fig. 4.**
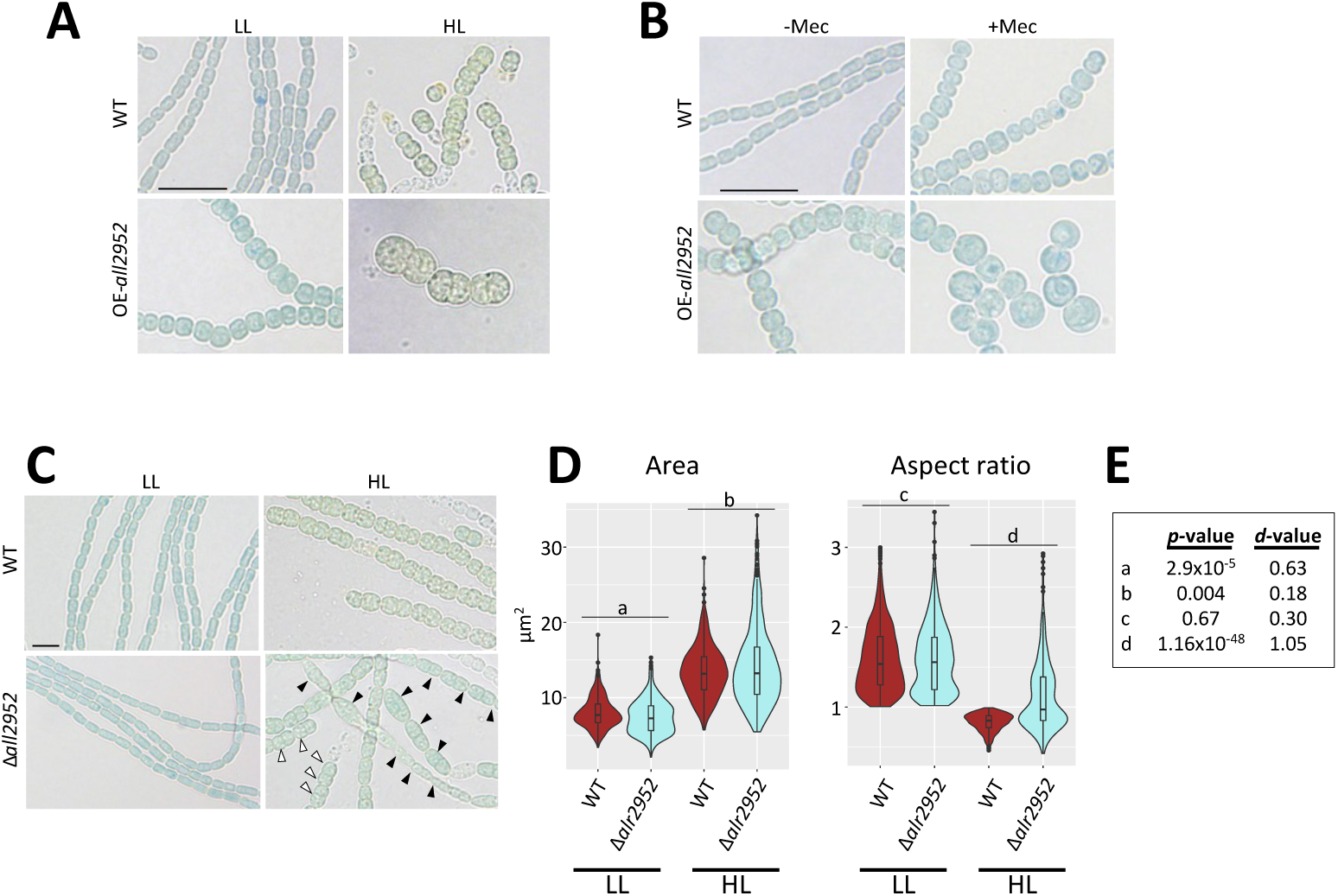
Impact of HL or mecillinam on OE-*all2952* cells and morphology of the Δ*all2952* mutant. (A) Pictures show light microscopy images of WT *Anabaena* or the OE-*all2952* strain incubated in LL or HL. All pictures were taken at the same magnification (scale bar, 10 µm). (B) Pictures of WT *Anabaena* or the OE-*all2952* strain cells incubated in the absence or presence of 40 µg ml-1 mecillinam. All pictures were taken at the same magnification (scale bar, 10 µm). (C) Brightfield microscopy images showing the phenotype of the WT and the Δ*all2952* mutant under LL (left panels) or HL (right panels). Empty and solid arrowheads point at cells with globular and non-globular morphology, respectively. All pictures were taken at the same magnification, scale bar 5 µm. (D) Violin plots of the area and aspect ratio of WT and Δ*all2952* cells cultured in LL or HL (*n*>490 cells). (E) Number indicate the *p-*value of two-tailed Welch’s-*t* test and the Cohen’s *d-*value of the pairwise comparisons indicated in D.

To assess the contribution of All2952 to the morphological change, a knock out mutant (Δ*all2952*) was generated (Fig. S8). While in LL this mutant was morphologically indistinguishable from the WT, HL cultures exhibited a heterogeneous mixture of elongated cells with a variety of sizes and shapes and some globular cells (Fig. 4C,D,E). It is worth mentioning that the ratio of elongated vs. globular cells was inconsistent in distinct replica, showing wide variations. PCR assays confirmed the uniformity of the Δ*all2952* genotype of the population in HL despite the heterogenous morphology. This observation suggested that while All2952 is important for achieving the large globular morphology in HL, in its absence, other aPBPs appear able to act as surrogates, enabling some cells of the population to become globular.

### Impact of cell morphology on light absorption and photosynthesis

An interesting finding was that under HL, WT cells showed a distinct distribution of thylakoids, which could be visualized via monitoring of the red fluorescence of chlorophyll *a* (Chl a) using confocal microscopy. Under LL, WT cells displayed red fluorescence mostly in the cytoplasm periphery, the central part being mostly devoid of fluorescence. In HL, fluorescence was at the periphery but also projected in the form of tangled or convoluted threads to the central part of the cytoplasm (Fig. 5A and Fig. S9A). Interestingly, cells of the OE-*all2952* strain under LL not only recapitulated the large globular morphology of the WT in HL, but also showed a strikingly similar distribution of fluorescence (Fig. 5A and Fig. S9A). To get a deeper insight, cells were analyzed by transmission electron microscopy (TEM). Noticeably, the positioning of thylakoids was in all cases consistent to the observed distribution of fluorescence (Fig. 5A, B). In LL, WT cells displayed in the periphery of the cytoplasm arrays of 4-6 thylakoid layers roughly running in parallel to the plasma membrane, the central part of the cell being virtually devoid of thylakoids. In HL, the number of thylakoid layers in the periphery decreased to 1-4 and most cells showed arrays of 1-4 layers in the center. Cells of the OE-*all2952* strain, showed a distribution of thylakoids similar to the WT in HL, with peripheral and central arrays; however, both peripheral and central layers were comparable in number to that of the WT in LL (Fig. 5B and Fig. S9B). As a whole, these results suggested that the overexpression of *all2952* not only induced a large globular morphology but also impelled thylakoids to invade the central part of the cytoplasm. In contrast, the reduction in the number of thylakoid layers was specifically induced by HL.

**Fig. 5.**
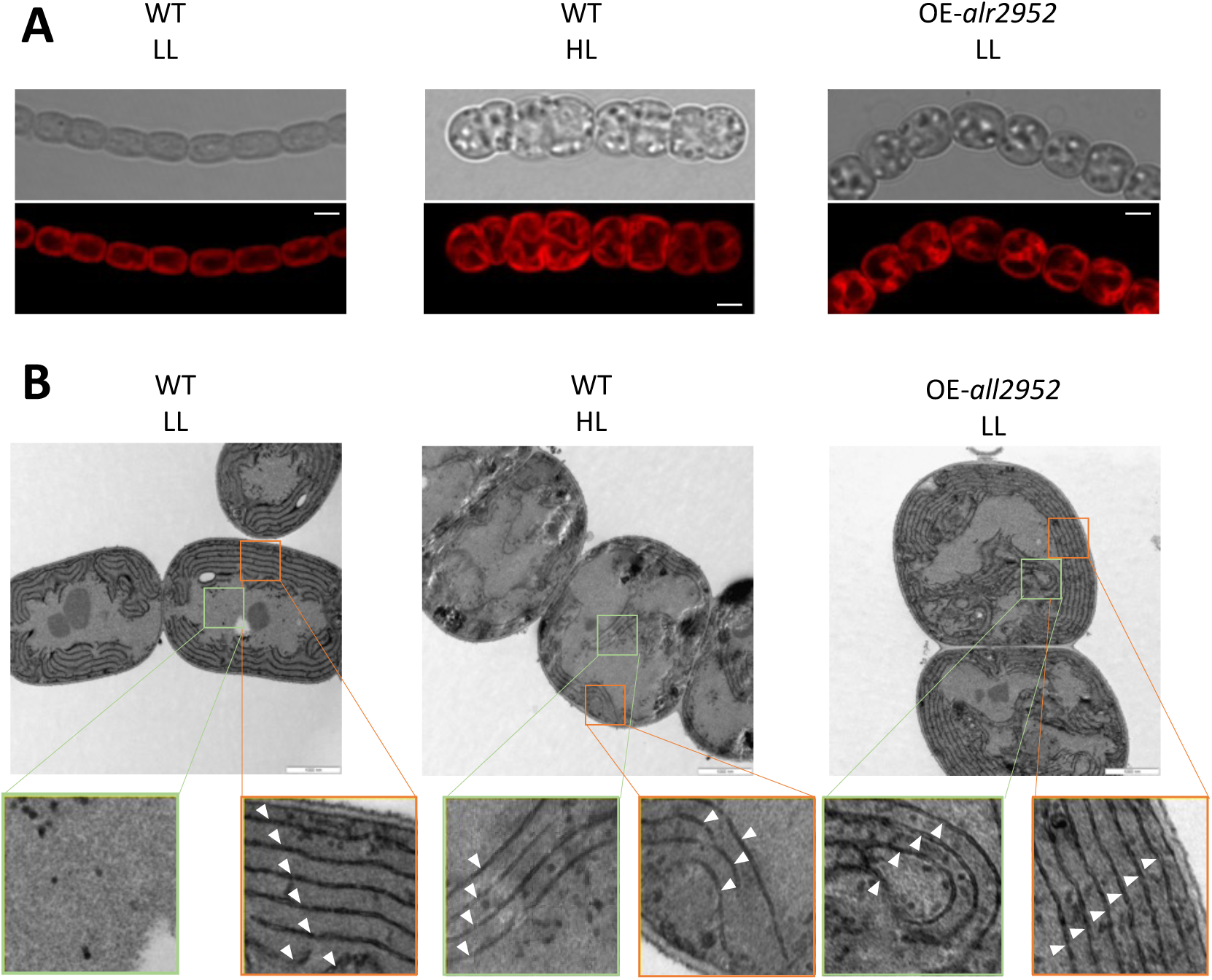
Fluorescence distribution and ultrastructure of *Anabaena* WT and the OE-*all2952* strain. (A) Cultures of WT *Anabaena* cultured in LL (left panels) or HL (middle panels), or the OE-*all2952* strain cultured in LL (right panels) were observed by confocal microscopy (Scale bar, 2 µm). Top panels are brightfield images, bottom panels are fluorescent images taken with excitation wavelength of 488 nm and emission wavelengths from 650-750 nm, so that the signal mostly corresponds to the fluorescence of Chl *a*. (B) Transmission electron microscopy images of the strains shown in (A). Cells were fixed with glutaraldehyde, stained with permanganate and thin sections were examined (Scale bar, 1 µm). Peripheral (orange frame) and central regions (green frame) were zoomed in to the same scale and are shown at the bottom. Arrowheads indicate thylakoid layers.

The morphological convergence of LL-grown OE-*all2952* and HL-grown WT cells established this overexpression strain as a robust model to isolate the impact of globular morphology as a single variable, independent of other light-intensity acclimation mechanisms such as thylakoid membrane degradation. To assess the impact of morphology and thylakoid distribution on photosynthesis, we first determined the pigment content of WT and OE-*all2952* cultures growing in LL. Quantitative analysis of pigments extracted by solvent treatment or cell lysis confirmed that the content of Chl *a*, phycobiliproteins and carotenoids showed the same proportion in both strains. Hence, cell suspensions of WT and OE-*all2952* containing equal pigment content were prepared (Fig. 6A). Notably, the absorption spectra of OE-*all2952* cell suspensions were invariably lower than those of WT across the visible range, despite the identical pigment concentration (Fig. 6B, left). This suggested an optical effect, where pigments are less efficient at light harvesting when packaged within large globular cells with centralized thylakoids. Supporting this, when cells were disrupted via lysozyme treatment and sonication, the absorption spectra of the resulting extracts were identical (Fig. 6B, right), confirming that the reduced absorption is a property of the intact cellular architecture.

**Fig. 6.**
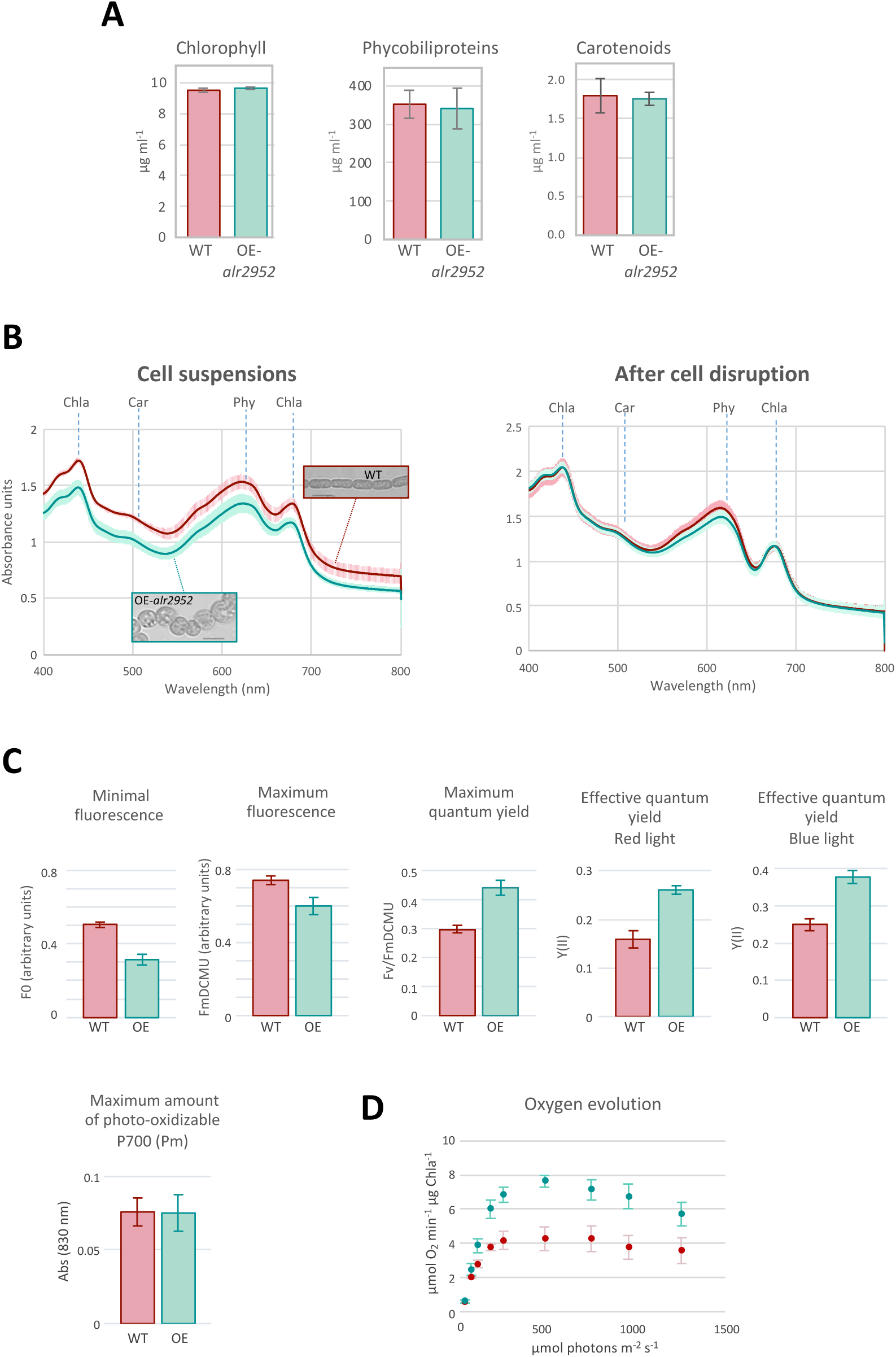
Pigment composition, absorbance and photosynthesis in *Anabaena* WT and the OE-*all2952* strain. (A) Cell suspensions containing 10 µg Chla ml-1 were prepared from cultures of *Anabaena* WT and the OE-*all2952* strain growing in LL. Pigments were extracted from the cell suspensions and quantitated. (B) Absorption spectra in the visible range of cell suspensions (left plot) prepared as in (A) or identical preparations after cell disruption by lysozyme treatment and sonication Inset pictures in the left plot correspond to confocal brightfield images (scale bar, 5 µm). (C) Plots show PAM measurements of WT and OE-*all2952* cell suspensions. (D) Oxygen production of WT and OE-*all2952* cell suspensions was measured in a Clark-type electrode at increasing light intensities as described in Materials and Methods.

We subsequently investigated the possible impact of cell morphology and thylakoid distribution on photosynthesis via Pulsed Amplitude Modulated (PAM) fluorometry. As shown in Fig 6C, the OE-*all2952* strain exhibited significantly lower minimal (*F*0) and maximal (*F*m) fluorescence than the WT. Since in cyanobacteria, these parameters integrate signals from both Chl *a* and phycobiliproteins, such decrease suggested an optical effect and was consistent with the observed loss in light absorption of OE-*all2952* cells. Despite this decrease in absolute fluorescence signals, the apparent maximum quantum efficiency (*F*v/*F*m) was consistently higher in OE-*all2952*, indicating that, while the globular cells absorb fewer photons per chlorophyll unit due to their optical properties, they are efficient at channeling the absorbed energy toward photochemistry. This was further corroborated by the higher effective quantum yield (Y(II)) recorded under both red and blue actinic light (Fig. 6C).

Further tests were performed to analyze the photo-oxidizable P700 signal (*P*m) via a shift in absorbance in the near-infrared, which is an indication of the amount of functional photosystem I (PSI). As shown in Fig. 6C, the PSI pool remained unchanged in the mutant, further supporting that the observed decrease of PSII fluorescence in OE-*all2952* stems from reduced excitation rather than reorganization of the photosynthetic complexes.

Finally, we assessed how the differential features of the OE-*all2952* strain could impact overall photosynthesis output upon short-term exposure (3 min) to light of non-saturating or saturating intensities. Oxygen evolution measurements evidenced that OE-*all2952* cells exhibited a robust activity at high irradiances (475 µmol photons m^-2^ s^-1^), while photosynthetic activity in WT cells saturated at moderate irradiances (240 µmol photons m^-2^ s^-1^) and showed signs of photoinhibition at higher light intensities (Fig. 6D). This suggested that the lower light-harvesting efficiency of globular cells confers a photoprotective advantage, limiting overexcitation and preventing photodamage, thereby enabling sustained photosynthetic performance under excess irradiance.

### Morphology of Anabaena in a simulated natural set

To get an insight into the role that cell morphology may play in natural populations, we considered the model of microbial mats. In nature, microbial mats form a few millimeters-deep layer of living matter over solid surfaces. Cyanobacteria are found frequently in intertidal mats and mats at the borders of freshwater bodies, but also in a variety of locations including desert crusts (Bolhuis *et al*., 2014; Perera *et al*., 2018). Although there are no records on the exact location where *Anabaena* sp. PCC 7120 was originally isolated, its metabolic capabilities indicate that this cyanobacterium likely dwells in microbial mats (Sarasa-Buisan *et al*., 2024; Olivan-Muro *et al*., 2025) as well as planktonically in freshwater. We set up an experimental system with artificial mats obtained by culturing *Anabaena* cells embedded in a semi-solid layer of BG11 medium supplemented with 1% agarose on top of BG11-agar medium (Figure 7A). Plates were illuminated for 48 hours with zenithal light of 100 µmol photons m^-2^ s^-1^ (termed “HL” in this experiment) or 20 µmol photons m^-2^ s^-1^ (LL) and cells from the top or the bottom of the cyanobacterial layer were observed by confocal microscopy. As shown in Fig. 7B, cells from the top of the artificial mat incubated in HL presented the typical HL phenotype described above, whereas those from the bottom had an elongated morphology resembling that of cells in LL. By contrast, cells from both the top and the bottom of the artificial mat incubated in LL had an elongated rod shape typical of LL (Fig. 7B). These observations bring into evidence that, within the same population, cells differentially adapt their morphology to the local incident light intensity.

**Fig. 7.**
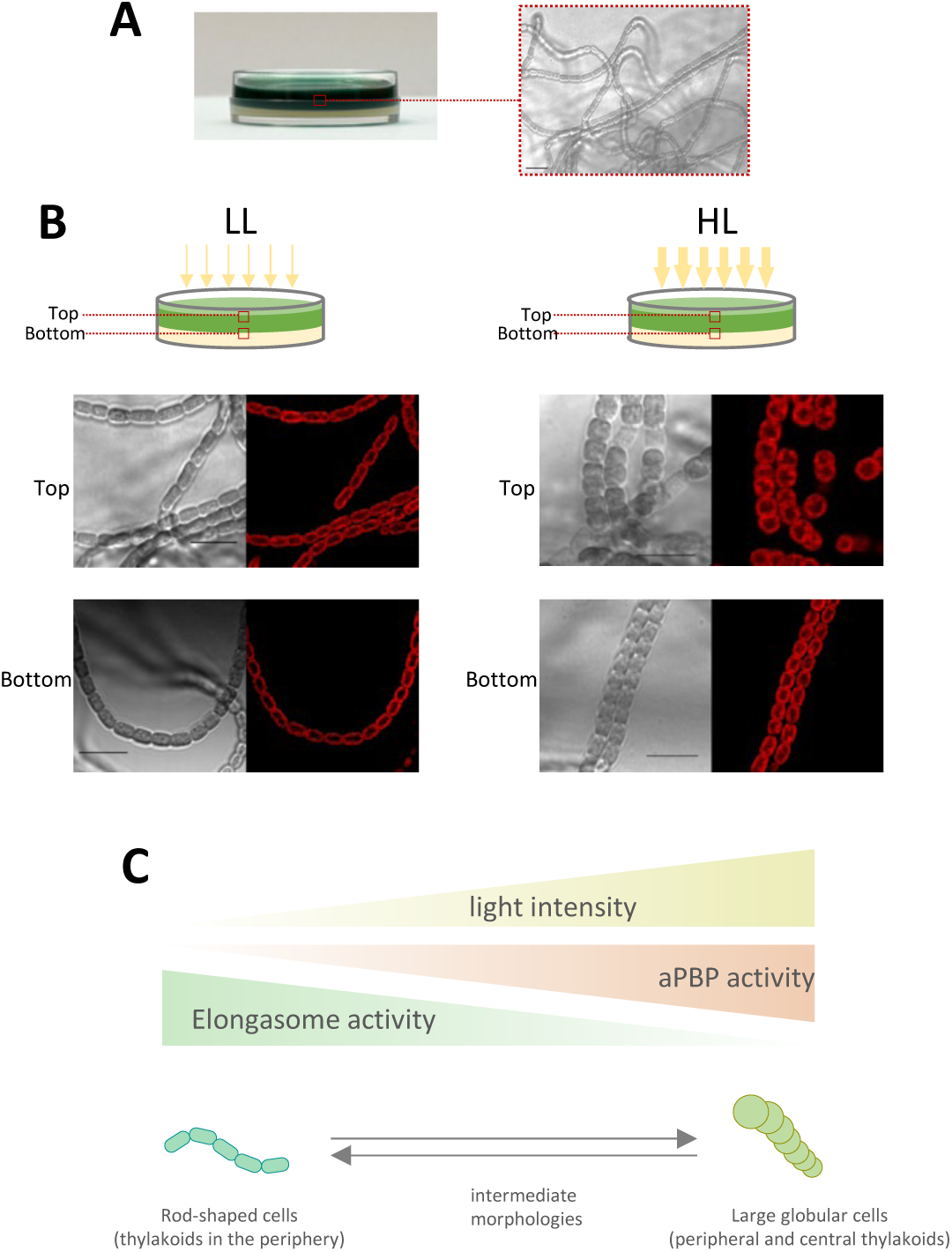
*Anabaena* morphology in artificial microbial mats. (A) Artificial microbial mats were prepared by embedding cells (50 µg Chla) in BG11 medium supplemented with 0.5% agarose on top of solid BG11-1.5% agar medium in 3.5 cm diameter Petri dishes. Left panel shows a representative image of a plate. The right panel is a brightfield confocal image showing *Anabena* filaments growing within the BG11-agarose matrix. Scale bar, 10 µm. (B) Plates prepared as in (A) were cultured in LL (20 µmol photons m-2 s-1) or in HL (100 µmol photons m-2 s-1) and sections from the top or the bottom of the cyanobacteria-containing layer were photographed by confocal microscopy (scale bar, 10 µm). Each panel shows brightfield confocal images (left) and fluorescence (excitation, 488 nm; emission 650-750 nm) images (right). (C) Diagram of the morphological changes of *Anabaena* as a function of light intensity. Light intensity and the activity of aPBPs and the elongasome are represented as triangles.

## Discussion

### The genuine morphology of Anabaena

*Anabaena* is commonly cultured in the laboratory with light intensities ranging from 20 to 60 µmol photons m^-2^ s^-1^. Consequently, most of our knowledge on this species pertains to cells growing under low or moderate irradiances. In this work we show that under HL intensities, *Anabaena* adopts a large globular morphology, distinct from the typical rod-shape in LL. The observation of intermediate morphologies at intensities between 20 and 200 µmol photons m^-2^ s^-1^, may explain the conflicting descriptions in previous reports, which defined *Anabaena* cells as rods, barrels or ellipsoids (Hu *et al*., 2007; Parsiegla *et al*., 2012; Velázquez-Suárez *et al*., 2022)

### Mechanisms underlying the transition into a large globular morphology

Evidence on model rod-shaped bacteria indicates that cell morphology is kept constant by balanced operation of PG assembly complexes, and deviations from this equilibrium cause morphological changes (Daniel *et al*., 2000; Formstone & Errington, 2005). In *Anabaena*, the elongasome is required for maintenance of the rod-shape in LL, defective mutants showing a less elongated shape (Burnat *et al*., 2014; Velázquez-Suárez *et al*., 2020), but it is dispensable for the large globular morphology in HL, achieved through the action of aPBPs. Despite this, it must be emphasized that elongasome Mre factors, particularly MreC, are required for proper growth in HL, bringing into evidence a novel unexpected function, apparently independent of known acclimation mechanisms (Fig S2, S3, File S1), which would be interesting to investigate.

Our observations indicate that in *Anabaena,* environmental conditions dictate which entity drives peripheral PG assembly; the elongasome being prevalent in LL, promoting a rod-shape, and aPBPs in HL, promoting a large globular morphology. The intermediate morphologies at moderate irradiances suggest progressive replacement of the former by the latter as light intensity increases (Fig. 7C). This aligns with observations in *B. subtilis* showing that artificial setting of elongasome and aPBP relative activities produce distinct morphologies, the former favouring thin elongated rod-shaped cells and the latter wider and rounder morphologies (Dion *et al*., 2019). This has been interpreted to result from the orientation of the PG chains assembled by each enzyme. Elongasome motion generates belt-like chains perpendicular to the cell long axis that constrict cell widening but allow elongation, whereas the short displacements of aPBPs in all directions drive assembly of PG chains with no preferred orientation, which also enables enlargement in width (Dion *et al*., 2019).

Our work sheds light on the mechanisms that establish the dominance of one enzymatic entity over the other and paves the way for future studies. Neither aPBP- nor elongasome subunit-encoding genes are transcriptionally regulated by light (Fig. S6). Instead, evidence indicates that the large globular morphology is achieved by post-transcriptional stimulation of aPBPs. In other bacterial systems, aPBP operation requires stimulation, most commonly via interaction with specific partners (Straume *et al*., 2021). For instance, in *E. coli*, PBP1a and PBP1b activity is stimulated by contact with outer membrane proteins LpoA and LpoB, respectively (Pazos & Vollmer, 2021). aPBP-activating partners in *Anabaena* are yet to be identified, and their possible interplay with light intensity remains to be explored.

Our evidence indicates that All2952 is the primary driver of the transition to a globular morphology under HL. All2952 activity in LL is likely minimal but not null. Consequently, in the OE-*all2952* overexpression strain, the cumulative effect of this basal activity is sufficient to induce a globular phenotype even in LL. Given that *all2952* expression is not induced by HL, we hypothesize that these conditions enhance All2952 activity through an as-yet unidentified mechanism (perhaps by interaction with a specific partner). This model is consistent with the observed phenotype of OE-*all2952* strain in HL, where stimulation of excess All2952 molecules likely drives massive peripheral PG synthesis, resulting in the giant globular morphology shown in Fig. 4A. Although peripheral PG synthesis in HL seems to be predominantly driven by All2952, the observation that a portion of cells in the Δ*all2952* mutant still become globular indicates partial functional redundancy of aPBPs in *Anabaena*, in line with reports in other species (Pazos & Vollmer, 2021; Straume *et al*., 2021)

Despite this partial redundancy, it is possible that each aPBPs has a specific function, becoming stimulated only in particular conditions. In line with this, three *Anabaena* aPBPs (All2981, Alr4579 and Alr5101) have been described to be required for nitrogen fixation (Lázaro *et al*., 2001; Leganés *et al*., 2005). Similarly, *E. coli* PBP1a and PBP1b display optimal activity at different pH range, which is thought to enable acclimation to distinct conditions (Mueller *et al*., 2019).

An important observation in this work is that the antibiotic mecillinam also induces a large globular morphology in *Anabaena* cells, suggesting that this drug influences cell-shape through mechanisms similar to HL. Consistently, mecillinam appears to stimulate the large aPBP load in the OE-*all2952* strain, inducing vast peripheral PG synthesis and a giant globular morphology (Fig. 4B). While mecillinam specifically inhibits PBP2 in *E. coli*, thereby blocking the transpeptidase activity of the elongasome (Cho *et al*., 2014), in *Anabaena* it induces a globular morphology also in the absence of PBP2 (Fig. 2). This indicates that in this species, mecillinam stimulates aPBP activity independently of other events.

Recent reports have unveiled new elements regulating cell size and/or shape in *Anabaena* or related strains (Sun *et al*., 2023; Zeng *et al*., 2023; Liu *et al*., 2024; Nguyen *et al*., 2025; Springstein *et al*., 2026). Interestingly, cyclic-di-GMP has been found to act as a second messenger that regulates cells size through a specific receptor (Sun *et al*., 2023; Zeng *et al*., 2023). Furthermore, RNA metabolism mediated by ribonuclease E/RebA also influences cell shape (Liu *et al*., 2024). It is also worth mentioning the strikingly similar morphology of mutants defective in a novel actin-like protein, named CorM in *Anabaena,* with the globular morphology shown in our study. (Nguyen *et al*., 2025; Springstein *et al*., 2026). While knowledge of the mechanisms governing cell morphology in *Anabaena* remains fragmentary, it would be interesting to investigate the interplay between these novel elements and their possible relation with light intensity.

### Structural and physiological features associated with the large globular morphology

Our data indicate that the size increase in HL is not driven by a necessity to accommodate rapidly produced biomass, as fast-growing, CO2-supplemented cultures maintain a normal size and rod shape (Fig. S1). Instead, the particular morphology observed in HL may be a requirement for reorganization of cytoplasmic content. In this regard, we observed in HL a rearrangement of thylakoids. HL-induced re-organization has also been documented in thylakoids of plant chloroplasts; for instance, in *Arabidopsis,* the grana disc diameter decreases while the gap between grana layers increases (Herbstová *et al*., 2012). This structural change facilitates membrane protein mobility, thereby enabling the repair of PSII photodamage. In the unicellular green alga *Chlamydomonas reinhardtii*, the number of thylakoid layers decreases and the luminal space increases, but the functional consequences are not well understood (Polukhina *et al*., 2016; Broderson *et al*., 2024; Dupuis *et al*., 2025). In *Anabaena,* we observed two distinct phenomena in HL. First, a reduction in the number of peripheral thylakoid layers, reflecting a decrease of the total thylakoid surface area. This likely curtails light absorption, which aligns with known photoacclimation mechanisms (Bailey & Grossman, 2008; Allahverdiyeva *et al*., 2015; Tiwari *et al*., 2025). Second, a re-positioning of thylakoids toward the center of the cytoplasm, which is inherent to the shift into a large globular morphology rather than a direct consequence of HL. Nevertheless, thylakoid redistribution would likely contribute to HL acclimation as well (see below). Notably, in the rod-shaped unicellular cyanobacterium *Synechococcus* sp. PCC 7942, while the number of thylakoid layers also decreases in HL, they do not protrude to the center of the cytoplasm and cells do not become globular (Huokko *et al*., 2021), further supporting a linkage between thylakoid centralization and the transition to a globular morphology.

The OE-*all2952* strain has provided a unique opportunity to uncouple shape-dependent events from light-induced stress responses. Evidence in this study showed that cell suspensions of this strain exhibit lower absorbance along the visible spectrum than WT cell suspensions with equal pigment content. This observation can be attributed to the *packaging effect*, an optical phenomenon describing the discrepancy in light absorption by pigments uniformly distributed in a solution versus their absorption when contained within discrete packages, such as cells (Duysens, 1956). Packaging reduces absorption efficiency, which is thought to be caused by self-shading of pigments. While packaging is influenced by many factors, theoretical and empirical work has demonstrated a correlation between this phenomenon and cell morphology; specifically, absorption is less efficient as cell size increases (Kirk, 1975, 1994; Morel & Bricaud, 1981). For decades, packaging has been acknowledged as a key phenomenon in large-scale studies on oceanic primary production, as it explains much of the discrepancy in Chl *a*-specific absorption recorded via satellite vs. *in situ* Chl *a* quantification upon extraction. Substantial correction has been achieved by accounting for the size of dominant phytoplankton species as a major cause of the packaging effect (Bricaud *et al*., 1995; Ciotti *et al*., 2002; Mouw & Yoder, 2010). Cell shape also impacts absorption; theoretical estimations predict that large spherical cells represent the most inefficient geometry for light collection, whereas thin cylindrical forms are much more effective (Kirk, 1976). Interestingly, thylakoid positioning has also been shown to contribute to the packaging effect (Berner *et al*., 1989).

While many studies on the packaging effect are blurred by the use of mixed populations with species of different morphologies and distinct pigment compositions, our work provides an ideal model where the cell size of a single species can be set by the control of a single enzyme. This enables the analysis of light absorption and photosynthesis as a function of morphology as the sole variable, isolated from other confounding factors. Our finding that cell disruption equalizes the absorption spectra of WT and OE-*all2952* (Fig. 6B) provides empirical evidence that the observed optical differences are purely structural.

Moreover, our results revealed pronounced effects of the large globular morphology and the associated thylakoid repositioning on photosynthesis. These effects are not apparently caused by a decrease or rearrangement of the photosynthetic complexes. Instead, the reduced light absorption of large globular cells seemingly modulates light harvesting by PSII, supporting sustained photosynthetic efficiency, likely by avoiding the over-reduction of the plastoquinone pool, the subsequent oversaturation of the electron transport chain, and because PSII is better protected from photodamage. This way of functioning allows large globular cells to maintain high photosynthetic output under irradiances that are photoinhibitory for rod-shaped cells. Taken together, these findings suggest that under conditions of excess light, a transition into a large globular morphology and centralization of thylakoids seemingly provides a photoprotective advantage by reducing light absorption.

### Morphological transitions as a dynamic acclimation mechanism

It is well known that some bacteria exhibit plasticity, transitioning between morphologies that often correlate with distinct environmental conditions, but only in a few cases the adaptive value of a specific morphology has been demonstrated (Yang *et al*., 2016). Notable examples include rod-shaped pathogens like *Legionella pneumophila* and uropathogenic *Escherichia coli*, which transition into filamentous forms to evade phagocytosis by immune cells (Justice *et al*., 2008). Likewise, certain marine bacteria develop into filaments exceeding 7 µm to avoid predation by eukaryotic protists, whose consumption is limited by the dimensions of their oral apparatus (Justice *et al*., 2008). Among cyanobacteria, morphological variation is also present. For instance, the helical-shaped genus *Arthrospira* exists as left- or right-handed spirals (Mühling *et al*., 2003) and it is worth to emphasize the pioneering work by B. Montgomery on the effect of light spectral quality on the morphology of *Fremyella diplosyphon* (Montgomery, 2015).

In this study, we show that *Anabaena* is a pleomorphic bacterium capable of transitioning between rod-shaped and large globular morphologies, each suited to specific light irradiances. Our results suggest that, in nature, *Anabaena* likely displays these distinct morphologies dynamically, depending on environmental variables, such as the season, its spatial position within a microbial mat, or its depth in the water column.

Cyanobacteria display numerous physiological and behavioral mechanisms for thriving in HL, including the excretion of protective sunscreen molecules, modulation of photosynthesis or the control of buoyancy via gas vesicles (Walsby *et al*., 1997; Tiwari *et al*., 2025). In light of the results presented here, we propose that the morphological shift undergone by *Anabaena* at the onset of HL should be regarded as a novel photoprotection mechanism, adding to the suite of strategies mediating HL acclimation.

## Acknowledgements

We are indebted to Alicia Orea (IBVF) and Juan Luis Ribas (Citius, US) for excellent technical assistance with microscopy. We are grateful to Jesús A. G. Ochoa de Alda for critical reading of the manuscript and helpful insights.

## Competing interests

Authors declare that they have no competing interests.

## Author contributions

Experimental work: CV-S, MJM-P, MAR, MB, IL

Data management and analysis: CV-S, MJM-P, MAR, LC-G

Contributed tools: RL-I

Contributed ideas: All authors

Funding: JLC, DJN, RL-I, LC-G, IL

Revised the manuscript: All authors

First draft of the manuscript: IL

## Data Availability

All underlying raw data supporting the figures and supplementary material of this manuscript have been deposited in Zenodo under DOI https://doi.org/10.5281/zenodo.20265913. To preserve the peer-review process, these data are temporarily under embargo. Due to manuscript length restructuring, some files have been repositioned; Table S2 provides a complete guide matching the figures in this final manuscript with their corresponding source folders in the Zenodo dataset.

Reviewers can access the dataset through the following link https://zenodo.org/records/20265914?preview=1&token=eyJhbGciOiJIUzUxMiIsImlhdCI6MTc3OTA4OTUzNCwiZXhwIjoxODAyNjQ5NTk5fQ.eyJpZCI6ImY5N2FjNDFlLWZkODctNDM1OS1hMTkwLTJkOGNmYzMwM2QyZCIsImRhdGEiOnt9LCJyYW5kb20iOiIwNGIyYjIwMDRmNDFmOWQ0OWQ5OTg4ODEwYzM4NDg0MyJ9.xpdfmru2iVssqhyr3C-PSKTYEiDYYd_uop9tCtnfihQ-_SoA8DbXgc4u8h7ohfer3kUXl5YalikQGt-S6aVVOA

## Funding

Grants PID2021-128477NB-I00 funded by MCIN/AEI/10.13039/501100011033/ FEDER, UE and PID2024-162446NB-I00 funded by MICIU/AEI/10.13039/501100011033/ FEDER, UE to IL; grant PID2021-123500NB-I00 funded by MCIN/AEI/10.13039/501100011033/ FEDER, UE to JLC; grant PID2023-146704O funded by MICIU/AEI /10.13039/501100011033/FEDER, UE to LCG; grant RYC2021-034768-I funded by MICIU/AEI/10.13039/501100011033 and NextGenerationEU/PRTR to RLI and grant Deutsche Forschungsgemeinschaft (DFG, Emmy Noether project award number NU 421/1) to D.J.N

## Supporting Information

**Fig. S1.** Light-driven morphological transitions are reversible and do not depend on the growth rate.

**Fig. S2.** Elongasome mutants in HL.

**Fig. S3.** Complementation of *mre* mutants.

**Fig. S4.** Growth of *mre* mutants in cultures supplemented with 1% CO2.

**Fig. S5.** Impact of mecillinam on *mre* mutant cell morphology.

**Fig. S6.** Expression of genes encoding proteins involved in peripheral PG synthesis.

**Fig. S7.** Impact of HL or mecillinam on OE-*all2952* cells.

**Fig. S8.** Construction and verification of mutant Δ*all2952*.

**Fig. S9.** Fluorescence distribution and ultrastructure of *Anabaena* WT and the OE-*all2952* strain.

**File S1.** Molecular basis for the phenotype of mre mutants in HL.

**File S2.** Table of correspondence of Figures with raw data deposited in Zenodo.

**Table S1.** Overexpression of genes encoding aPBPs.

**Table S2.** Oligonucleotides used in this work.

## Notes

https://doi.org/10.5281/zenodo.20265913

### Competing Interest Statement

The authors have declared no competing interest.

