## Supplementary material for "The cyanobacterium *Anabaena* uses pleomorphism as an acclimation strategy to high light stress": Fig. S1

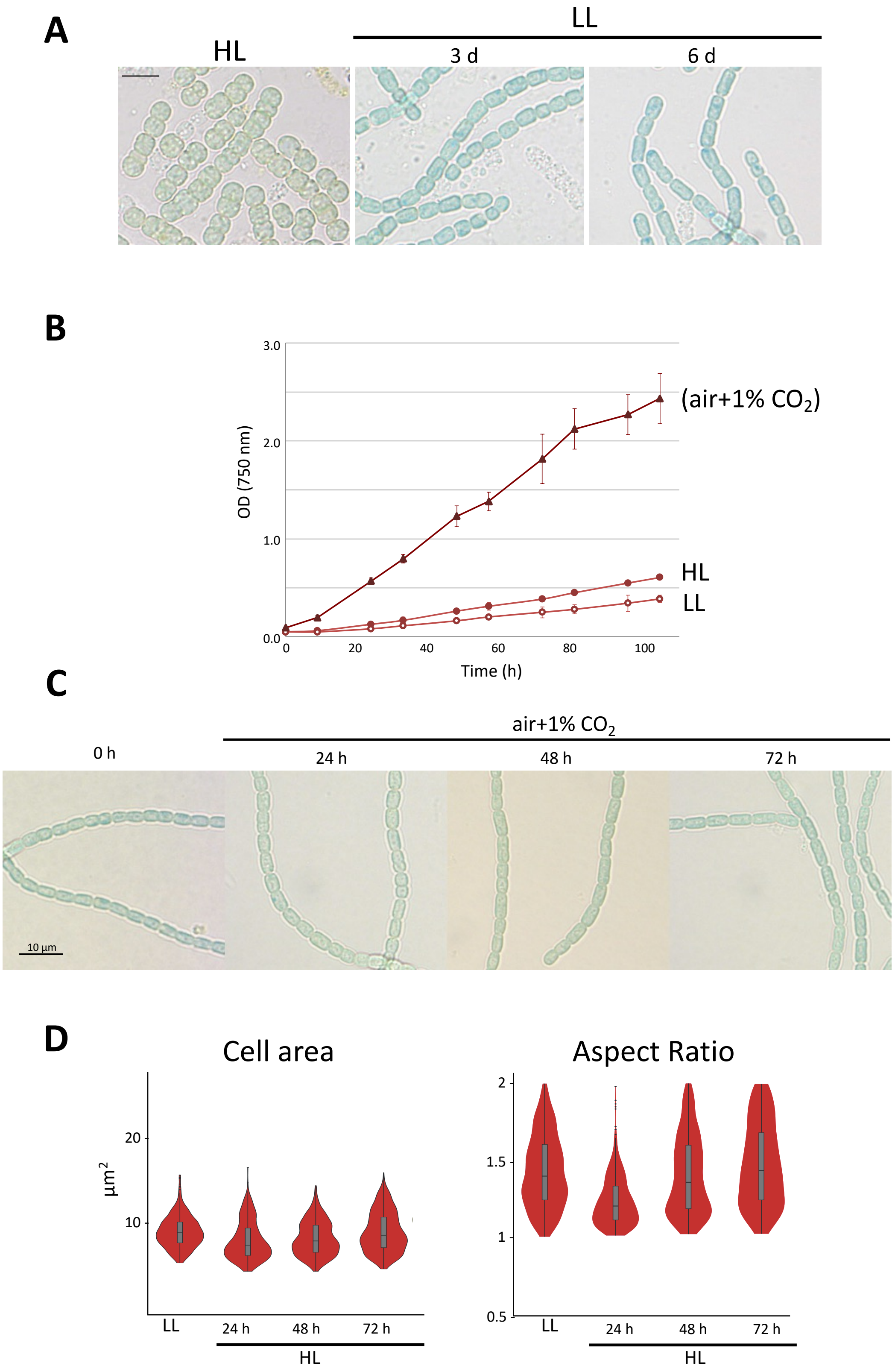


**Fig. S1. Light-driven morphological transitions are reversible and do not depend on the growth rate.** (A) *Anabaena* cells were cultured in HL (500 µmol photons m$^{-2}$ s$^{-1}$) for 72 h and transfered to LL (20 µmol photons m$^{-2}$ s$^{-1}$) for the time indicated in days (d). Scale bar 5 µm. (B) Growth curves of *Anabaena* liquid cultures incubated in LL, HL or in LL supplemented with 1% $CO_2$. (C) Brightfield microscopy images of cultures supplemented with 1% $CO_2$ at the indicated times. (D) Plots represent the cell area and the aspect ratio of cells from cultures supplemented with 1% $CO_2$ ($n$=500 cells).
