## Supplementary material for "The cyanobacterium *Anabaena* uses pleomorphism as an acclimation strategy to high light stress": Fig. S2

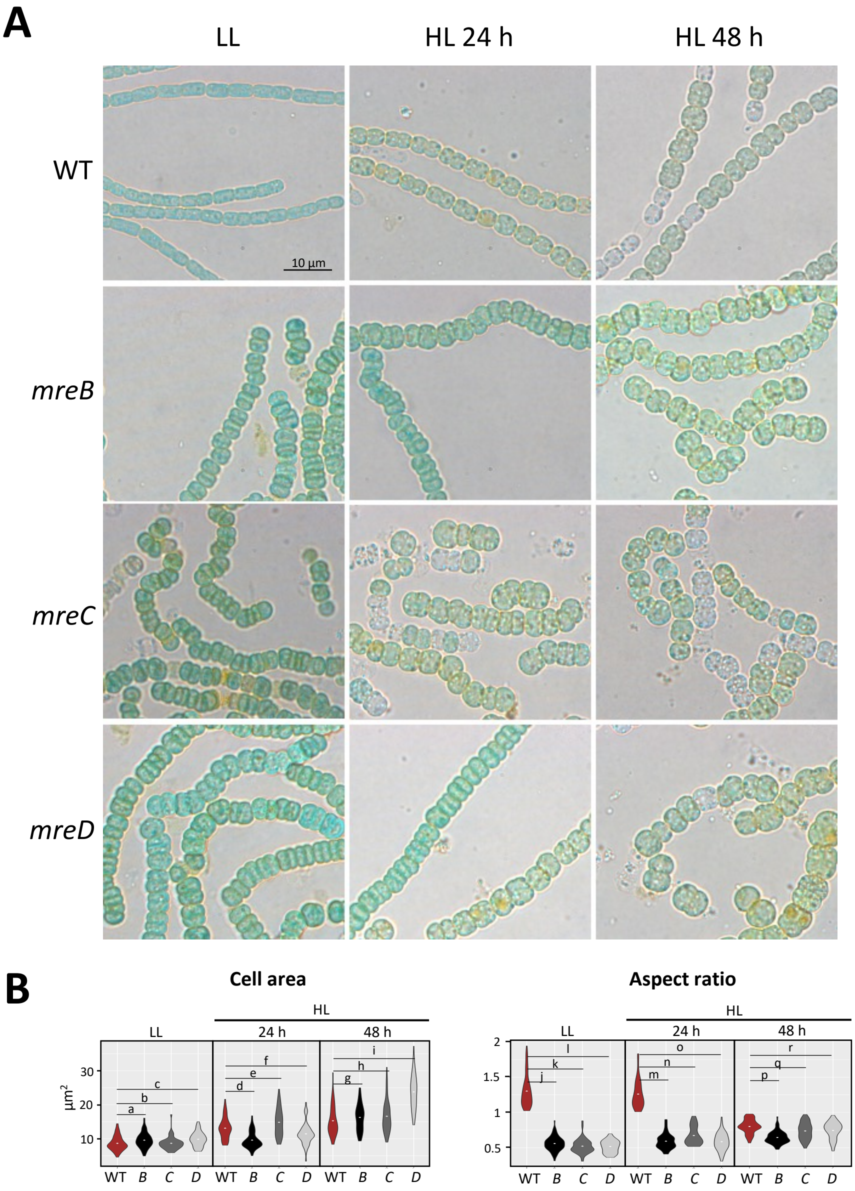


**C**

| | *p*-value | *d*-value | | *p*-value | *d*-value |
|---|---|---|---|---|---|
| **a** | 0.0002 | 0.30 | **j** | $4.8 \times 10^{-187}$ | 3.84 |
| **b** | 0.11 | 0.13 | **k** | $3.5 \times 10^{-188}$ | 3.84 |
| **c** | $3.5 \times 10^{-10}$ | 0.52 | **l** | $9.7 \times 10^{-193}$ | 4.46 |
| **d** | 0.0008 | 0.27 | **m** | $2.4 \times 10^{-89}$ | 1.99 |
| **e** | $5.2 \times 10^{-6}$ | 0.5 | **n** | $1.1 \times 10^{-22}$ | 1.14 |
| **f** | $7.9 \times 10^{-8}$ | 0.44 | **o** | $1.5 \times 10^{-61}$ | 1.56 |
| **g** | 0.98 | 0 | **p** | $7.9 \times 10^{-40}$ | 1.17 |
| **h** | 0.02 | 0.24 | **q** | $9.8 \times 10^{-6}$ | 0.47 |
| **i** | $3.7 \times 10^{-49}$ | 1.35 | **r** | $1.9 \times 10^{-7}$ | 0.43 |

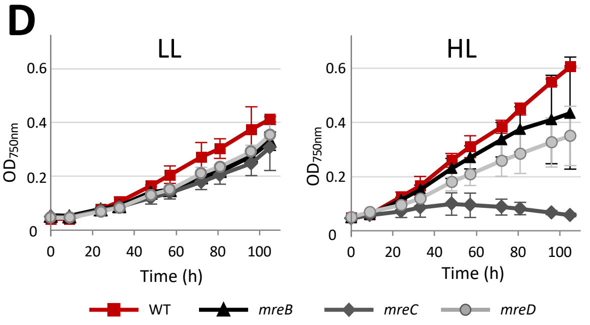


**Fig. S2.** Elongasome mutants in HL. (A) WT *Anabaena* and *mreB*, *mreC and mreD* mutants defective in elongasome subunits cultured in LL were transferred to HL and photographed under the light microscope at the indicated hours. All pictures were taken with the same magnification. (B) Rotated kernel density plots (as in Fig. 1) represent the cell area and the aspect ratio of cells from the experiment described in (A). The experiment was repeated 3 times and cells from all three experiments were quantitated (*n*=300 cells, except for the *mreC* mutant in HL (n=150 cells) due to viability loss). (C) Numbers correspond to the *p-value* of two-tailed Welch's-*t* test and the Cohen's *d* value for the indicated comparisons. Growth curves of *Anabaena* WT and the *mre* mutants in LL and HL. Data represent the average±SD of three independent experiments.
