## Supplementary material for "The cyanobacterium *Anabaena* uses pleomorphism as an acclimation strategy to high light stress": Fig. S3

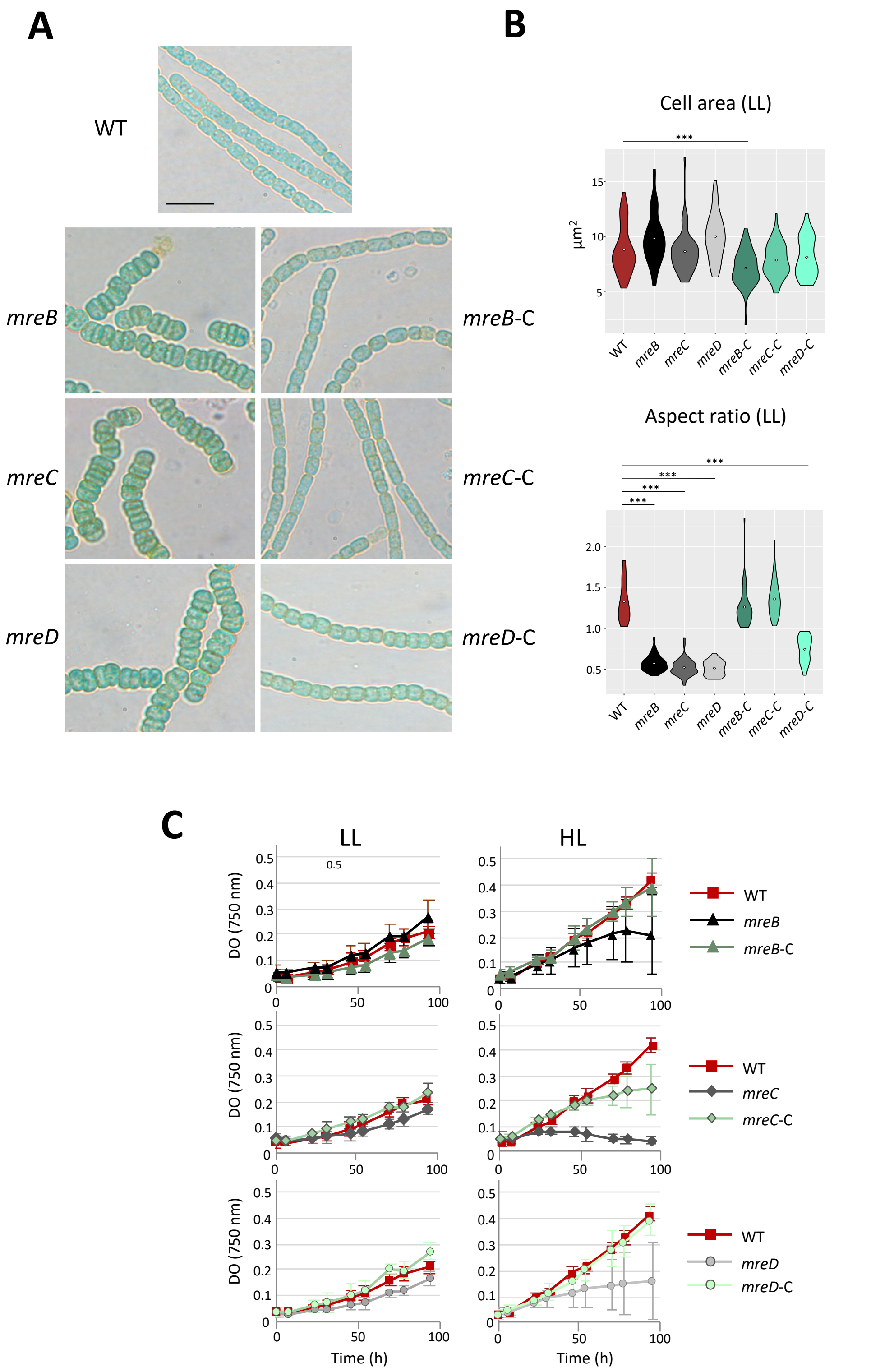


**Fig. S3.** Complementation of *mre* mutants. (A) Δ*mre* mutants were complemented as described in Materials and Methods and the morphology of the mutant (left) and complemented (right) strains growing in LL was compared. All pictures were taken to the same magnification (scale bar, 10 µm). (B) Kernel density plots represent the cell area and the aspect ratio of cells cultured as in A (*n*=50 cells). The significance of differences was tested using the Welch's-*t* test (*** means $p <$ 0.001). (C) Growth curves of liquid cultures of WT *Anabaena,* Δ*mre* mutants and complemented strains incubated in LL (left) or HL (right).
