## Supplementary material for "The cyanobacterium *Anabaena* uses pleomorphism as an acclimation strategy to high light stress": Fig. S4

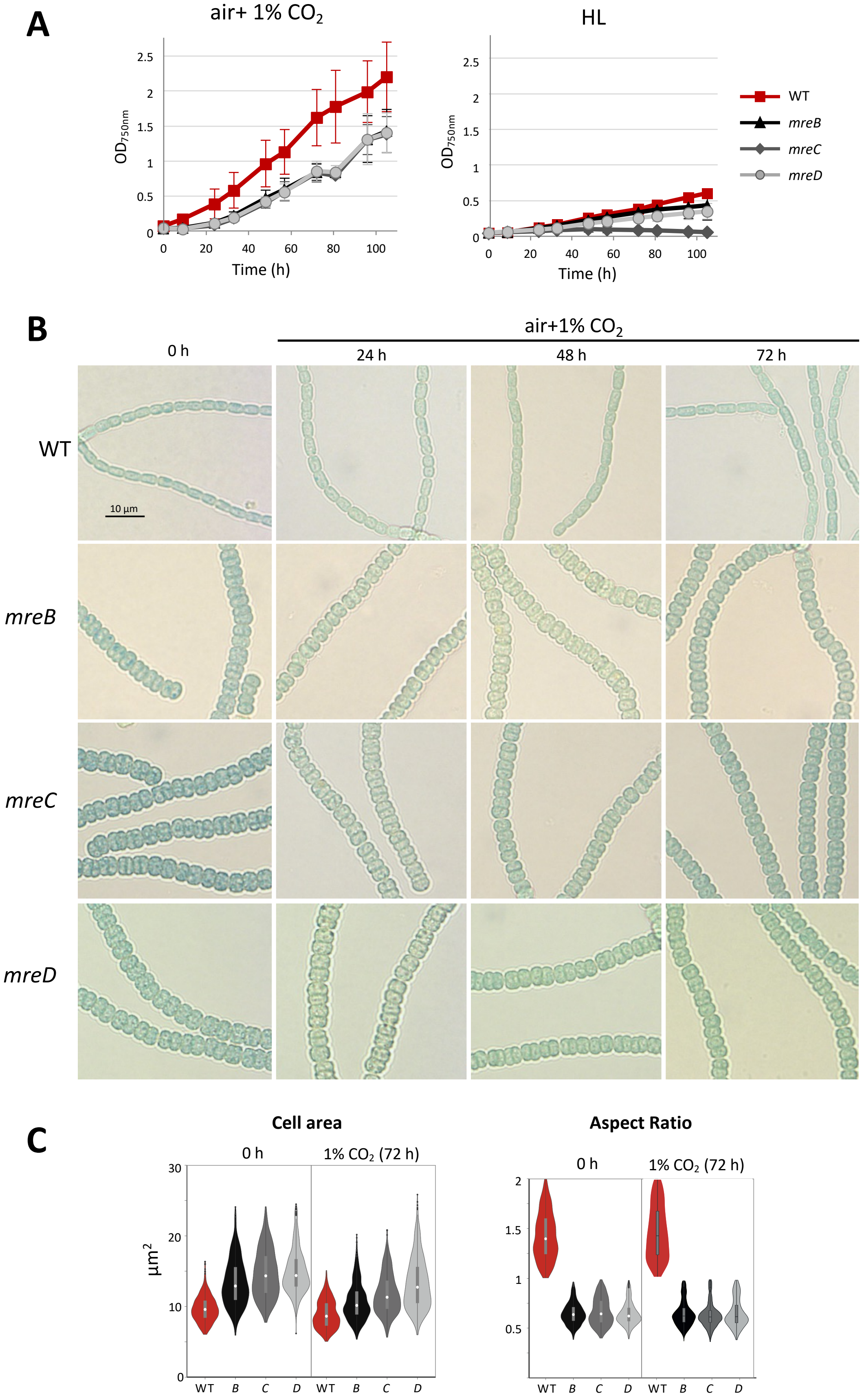


**Fig. S4.** Growth of *mre* mutants in cultures supplemented with 1% $CO_2$. (A) Growth curves of WT *Anabaena* and *mre* mutants growing in liquid cultures supplemented with 1% $CO_2$ (left). For comparison, growth curves of *Anabaena* WT and *mre* mutants growing in HL are provided. (B) WT *Anabaena* and *mre* mutants cultured in LL were incubated with $CO_2$ enrichment and photographed at the indicated times. (C) Plots represent the cell area and the aspect ratio of cells cultured as in B for the indicated time (*n*=500 cells).
