## Supplementary material for "The cyanobacterium *Anabaena* uses pleomorphism as an acclimation strategy to high light stress": Fig. S5

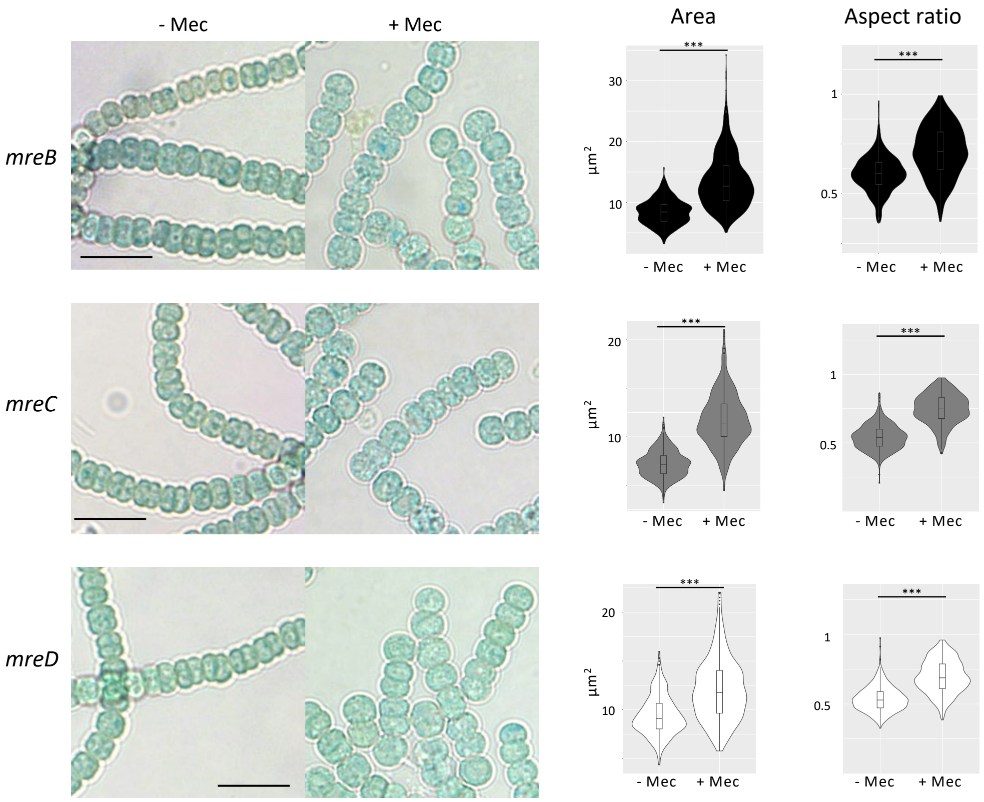


**Fig. S5.** Impact of mecillinam on *mre* mutant cell morphology. Cultures of *mre* mutants containing 1 µg Chla ml$^{-1}$ were supplemented or not with 40 µg ml$^{-1}$ mecillinam, as indicated, incubated for 72 hours in LL and observed by light microscopy. All pictures were taken with the same magnification (Scale bar 10 µm). Diagrams at the left side represent the cell area and the aspect ratio of cells from this experiment. The experiment was repeated three times and cells from the three experimental replica were quantitated ($n$=500). Results were statistically analized by Welch's-$t$ test (***, $p$<0.001).
