## Supplementary material for "The cyanobacterium *Anabaena* uses pleomorphism as an acclimation strategy to high light stress": Fig. S6

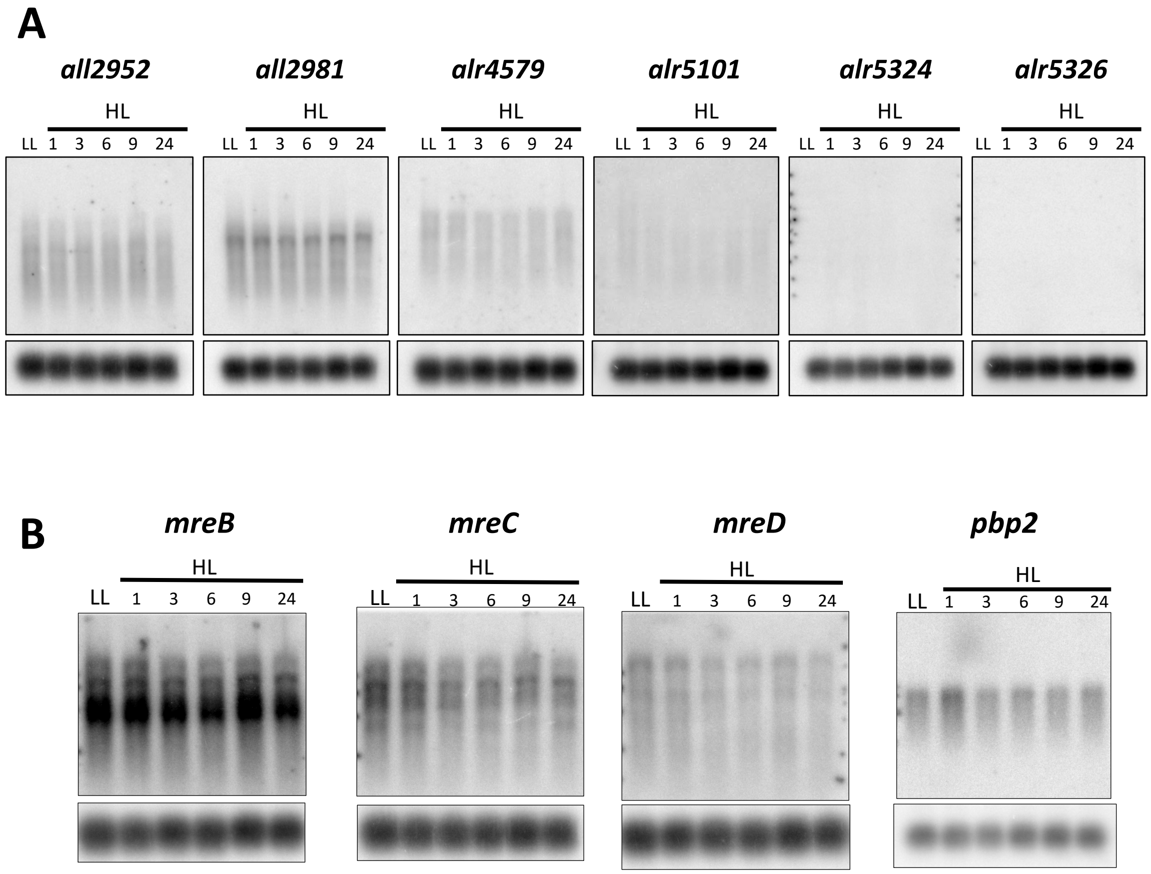


**Fig. S6.** Expression of genes encoding proteins involved in peripheral PG synthesis. (A) 5 µg of RNA isolated from cells grown in LL or incubated in HL for the time indicated (in hours) was loaded on a 1% agarose gel, transferred to a nylon membrane and hybridized to specific probes of aPBP-encoding genes. Each membrane was thereafter hybridized with a specific probe of the gene encoding the 5S rRNA as a loading control. (B) Details are like in A, but membranes were hybridized with probes of the indicated elongasome factor-encoding genes.
