## Supplementary material for "The cyanobacterium *Anabaena* uses pleomorphism as an acclimation strategy to high light stress": Fig. S7

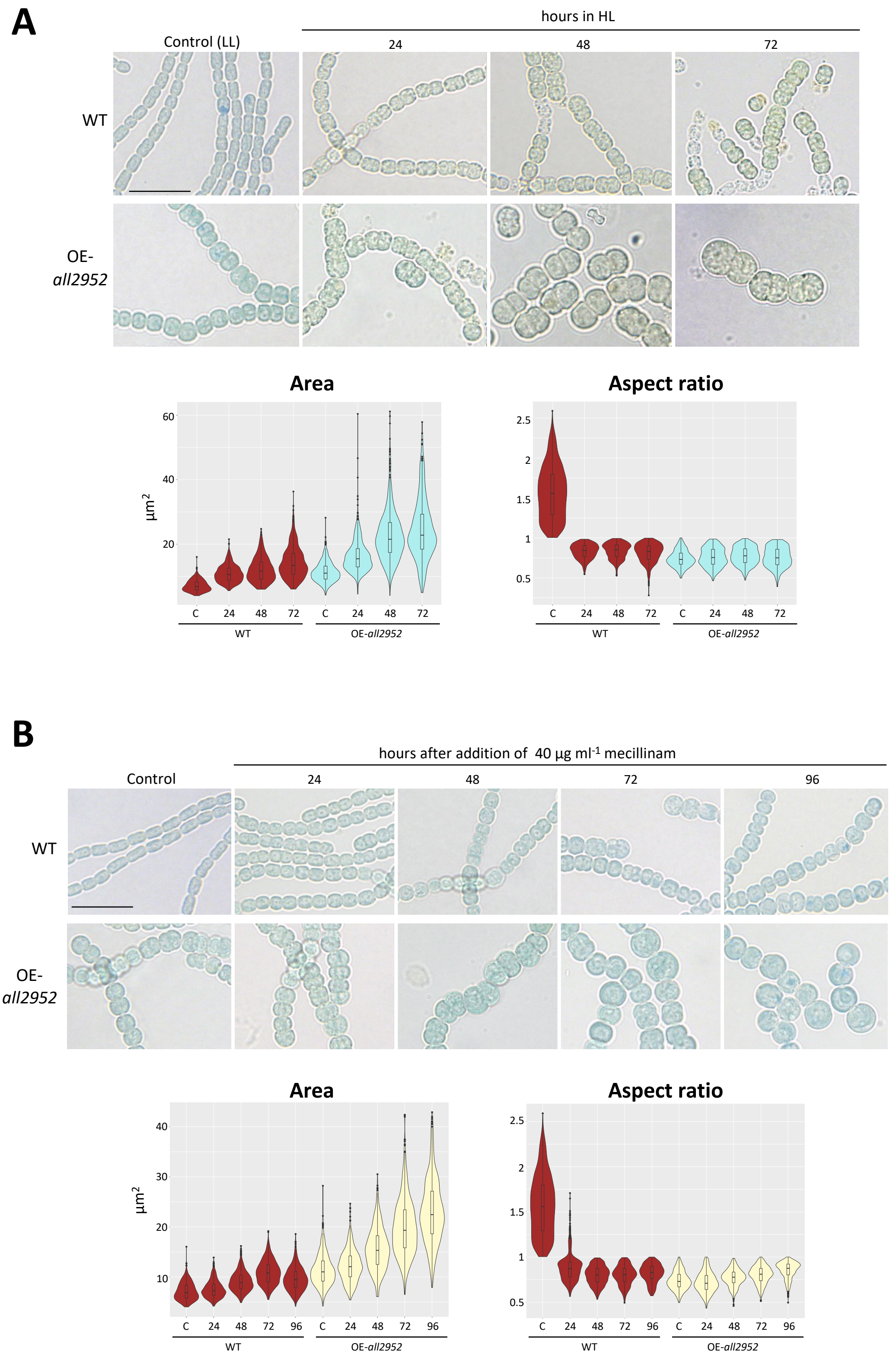


**Fig. S7.** Impact of HL or mecillinam on OE-*all2952* cells. (A) Cultures of WT *Anabaena* or the OE-*all2952* strain incubated in LL (c, control) were transferred to HL for the indicated time (in hours) and observed by light microscopy (scale bar, 10 µm). All pictures were taken at the same magnification. Diagrams represent the cell area and the aspect ratio of the indicated strains (the experiment was repeated three times, at least 400 cells of each timepoint were quantitated). (B) Cultures of WT *Anabaena* or the OE-*all2952* strain cultured in LL (c, control) were supplemented with mecillinam to a final concentration of 40 µg ml$^{-1}$ for the indicated time (in hours) and observed by light microscopy (scale bar, 10 µm). All pictures were taken at the same magnification. Diagrams represent the cell area and the aspect ratio of the indicated strains (the experiment was repeated three times, 500 cells of each timepoint were quantitated).
