## Supplementary material for "The cyanobacterium *Anabaena* uses pleomorphism as an acclimation strategy to high light stress": Fig. S8

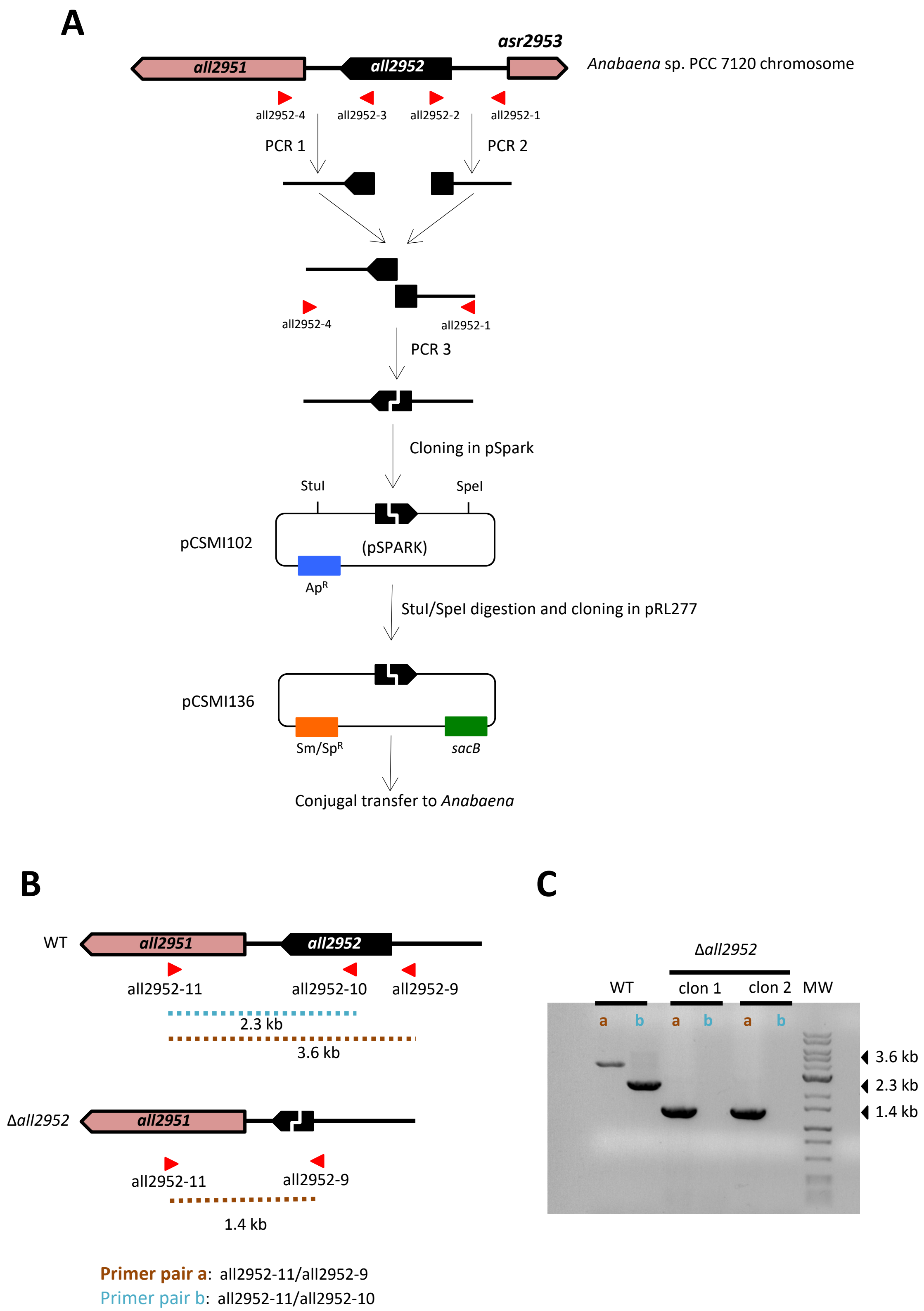


**Fig. S8.** Construction and verification of mutant Δ*all2952*. (A) Construction scheme of Δ*all2952*. Primers used for PCR are depicted as red arrowheads and are listed in Suppl. Table 2. Other details are indicated in Materials and Methods. (B) Scheme of the *all2952* genomic region in *Anabaena* WT and the Δ*all2952* mutant. Primer pairs used for verification of the genomic structure of the *all2952* locus and the expected size of the PCR products are shown. (C) Agarose gel with PCR products obtained using as template genomic DNA from *Anabaena* WT or from two different clones of the Δ*all2952* mutant. Primer pairs used for PCR are indicated in (B). MW, molecular DNA ladder. The size of PCR fragments is indicated at the left.
