## Supplementary material for "The cyanobacterium *Anabaena* uses pleomorphism as an acclimation strategy to high light stress": Fig. S9

A

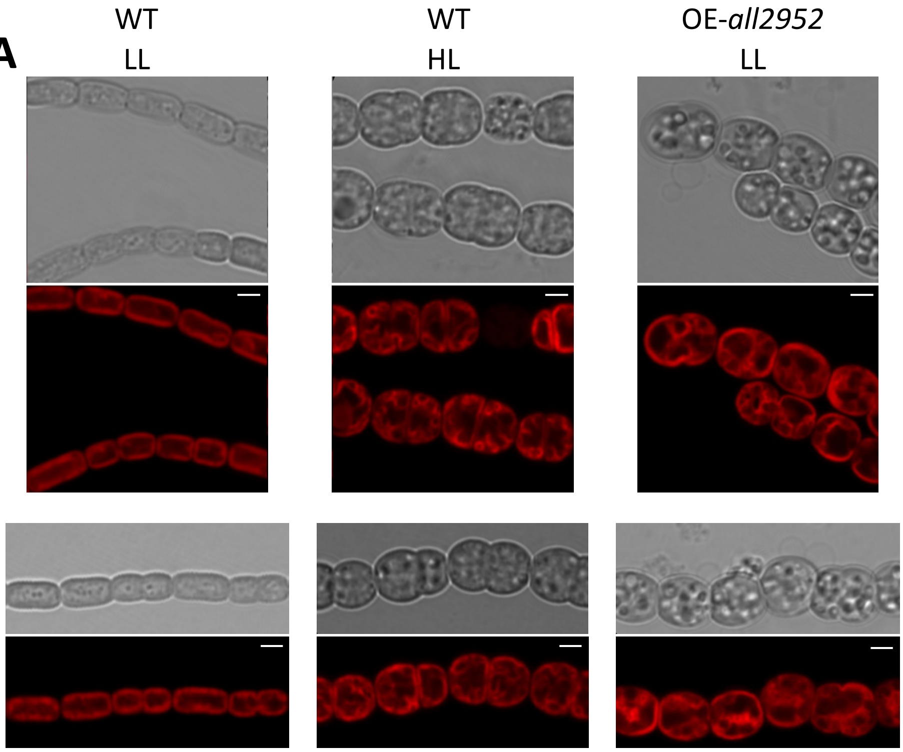


B

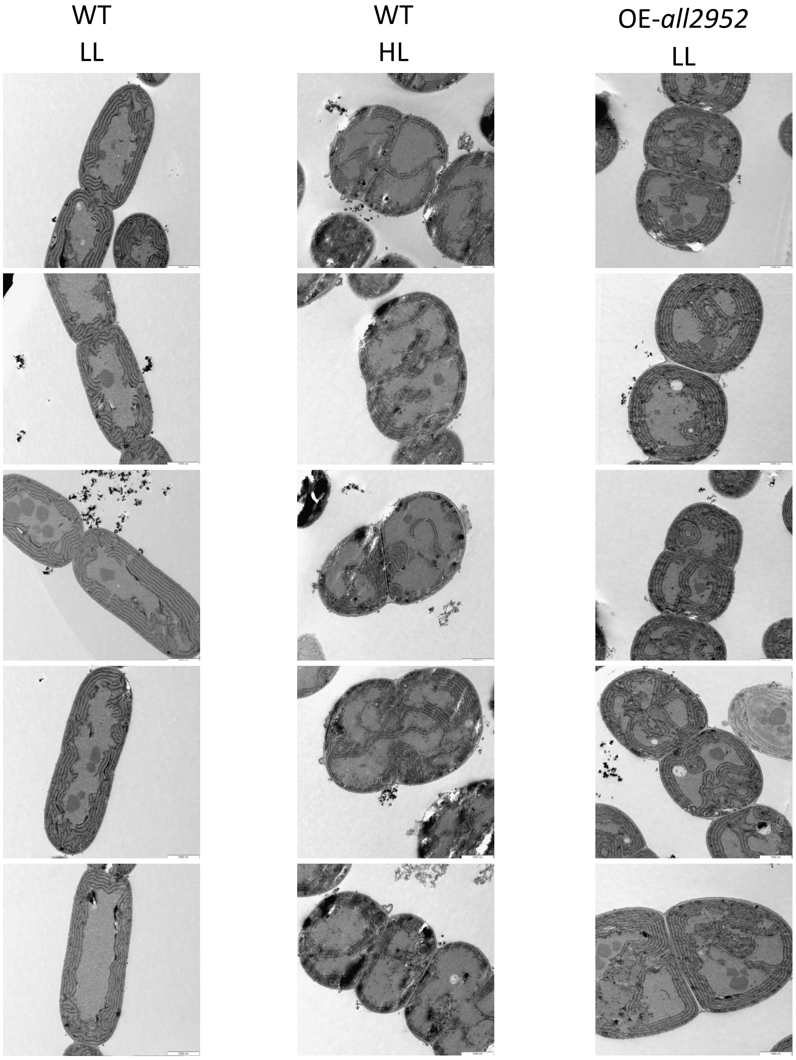


**Fig. S9. Fluorescence distribution and ultrastructure of *Anabaena* WT and the OE-*all2952* strain.** (A) Cultures of WT *Anabaena* cultured in LL (left panels) or HL (middle panels), or the OE-*all2952* strain cultured in LL (right panels) were observed by confocal microscopy (Scale bar, 2 μm). Top panels are brightfield images, bottom panels are fluorescent images taken with excitation wavelength 488 nm and emission wavelength 650-750, so that the signal mostly corresponds to the fluorescence of Chla. (B) Transmission electron microscopy images of the strains shown in (A). Cells were fixed with glutaraldehide, included, stained with permanganate and thin sections were photographed (Scale bar, 1 μm).
