## Supplementary material for "The cyanobacterium *Anabaena* uses pleomorphism as an acclimation strategy to high light stress": File S1

### *Molecular basis for the phenotype of mre mutants in HL*

As described above, each *mre* mutant showed two phenotypic alterations with no apparent relation namely, a discoidal morphology in LL and an impairment for growth in HL, which is particularly severe for the *mreC* mutant. Growth in HL requires a thorough acclimation of the photosynthetic apparatus and failure to initiate these mechanisms is detrimental for growth and survival (REF). Several approaches were undertaken to analyze whether these mechanisms functioned properly in the *mre* mutants:

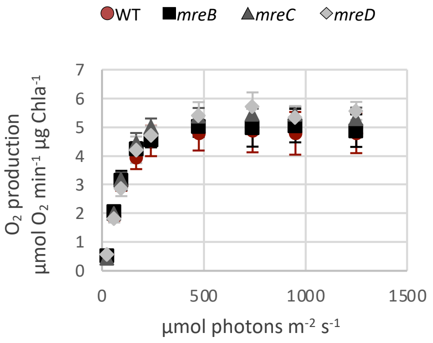


**Plot 1. Short-term response to increasing light intensities**

The overall functioning of photosynthesis at different light intensities was tested in a Clark-type electrode. Cell suspensions were exposed for 3 minutes to non-saturating or saturating light intensities (up to 1250 µmol photons $m^{-2}$ $s^{-1}$). As shown, the oxygen production rate of the wild type and the three *mre* mutants was similar.

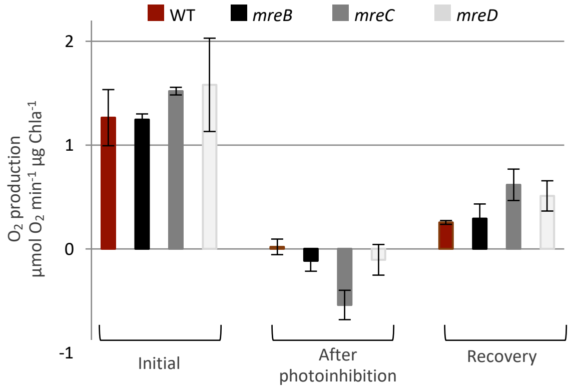


**Plot 2. Recovery of PSII after photoinhibition**

An important issue of the acclimation response is the recovery of photosystem II (PSII) from oxidative damage induced by HL, which inhibits photosynthetic activity unless repair mechanisms are induced. We analyzed oxygen production in a Clark-type electrode using suspensions of cells cultured in LL. After initial recording of the $O_2$ production (Initial), suspensions were exposed to the very high irradiance (1400 µmol photons $m^{-2}$ $s^{-1}$) for 1h and monitored (After photoinhibition). Absence of oxygen production by WT and *mre* mutants revealed PSII inactivation by photodamage. Thereafter, suspensions were incubated for an additional hour under low irradiance conditions (20 µmol photons $m^{-2}$ $s^{-1}$) and monitored again. As shown, $O_2$ production was restored for the wild type and the mutants to a similar extent (Recovery), indicating that PSII repair mechanisms were intact in the *mre* mutants.

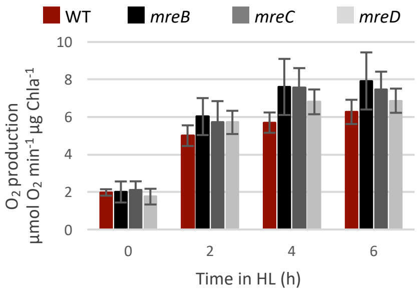


**Plot 3. $O_2$ production during the transition from LL to HL**

Cells cultured in LL (20 µmol photons $m^{-2}$ $s^{-1}$) were transferred to HL, (500 µmol photons $m^{-2}$ $s^{-1}$). $O_2$ production was measured before transfer (0 h), or after 2, 4 and 6 h of growth in HL.

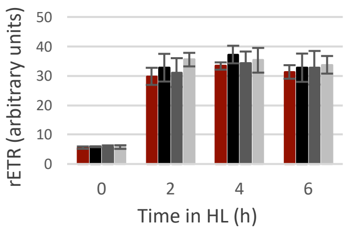


**Plot 4. Photosynthetic electron transport rate (rETR) during the transition from LL to HL**

Cells were cultured as above and rETR was measured before transfer (0 h), or after 2, 4 and 6 h of growth in HL by PAM. The relative ETR (rETR) was calculated as rETR = ΦPSII × PAR × 0.5, where ΦPSII is the PSII quantum yield, PAR is the incident photosynthetically active radiation, and 0.5 accounts for the fraction of light absorbed by PSII [2].

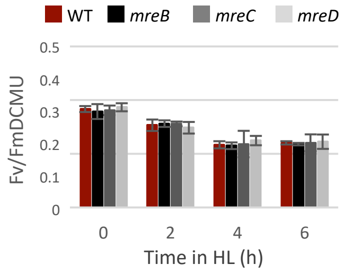


**Plot 5. Quantum yield kinetics during the transition from LL to HL**
Cells were cultured as above and Fv/Fm in the presence of DCMU was measured before transfer (0 h), or after 2, 4 and 6 h of growth in HL by PAM.

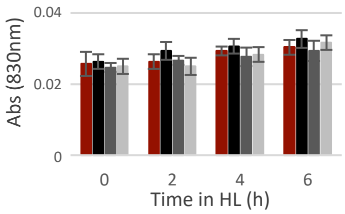


**Plot 6. Maximum photo-oxidatable $P_{700}$ during the transition from LL to HL**
Cells were cultured as above and the maximum photo-oxidatable $P_{700}$ (Pm) was measured before transfer (0 h), or after 2, 4 and 6 h of growth in HL by PAM

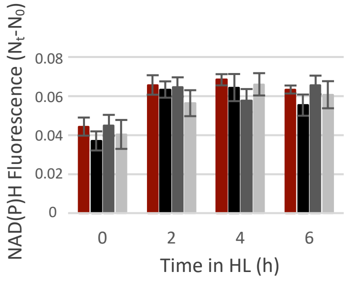


**Plot 7. NADPH Fluorescence during the transition from LL to HL**
Cells were cultured as above and the NADPH Fluorescence was measured before transfer (0 h), or after 2, 4 and 6 h of growth in HL by PAM. NAD(P)H fluorescence was measured using the NADPH/9-AA module of the DUAL-PAM-100, and was calculated as the difference between $N_t$, the steady-state NAD(P)H fluorescence during actinic illumination (10 min), and $N_0$, the basal NAD(P)H fluorescence [2].

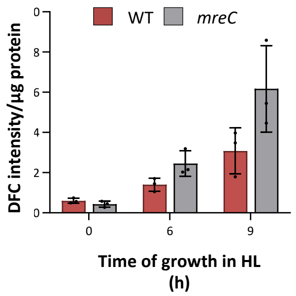


**Plot 8. ROS content of WT and *mreC* cells in HL**
WT and *mreC* cell suspensions were cultured for the indicated time (in hours) in HL. ROS levels were determined as previously described [2]. 5-(and-6)-chloromethyl-2',7'-dichlorodihydrofluorescein diacetate (CM-H2DCFDA) was added to a final concentration of 25 µM to the samples. After incubation for 30 min at 30°C, fluorescence (485 nm excitation wavelength and 525 nm emission wavelength) was measured in a Varioskan LUX fluorometer (Thermo Scientific).

### Conclusions

According to our observations above, the photosynthetic activity of the three *mre* mutants showed a similar short-term response to increasing light intensities as the WT, with similar $O_2$ production rates at all intensities and signs of photoinhibition at about 200 µmol photons $m^{-2}$ $s^{-1}$ (Plot 1). *mre* mutants also showed similar recovery of PSII after photodamage, indicating that PSII photorepair mechanisms are intact in these mutants (Plot 2). As expected, the WT showed an increase of the $O_2$ production rate and the rETR during the first 6 hours in HL and similar increases were observed for the *mre* mutants (Plots 3 and 4). Fv/FmDCMU declined similarly for the WT and the *mre* mutants, evidencing a similar decrease in PSII activity due to photodamage (Plot 5). The Pm increased for the WT and the *mre* mutants in a similar way, likely reflecting an increased capacity to prevent over-reduction of the electron transport chain under high light (Plot 6). Furthermore, the NADPH fluorescence increased similarly for all strains during the first 6 hours in HL (Plot 7), which is suggestive of some saturation of the CBB cycle, but with no signs of decoupling of light and dark reactions of photosynthesis. As a whole, these observations indicate that photosynthetic performance, as well as short- and mid-term acclimation mechanisms to HL, are functional in the *mre* mutants.

In an effort to investigate the basis for the lethal phenotype of the *mreC* mutant in HL, ROS accumulation was measured. As shown in Plot 8, after 9 h in HL, this mutant accumulated 2-fold the content of ROS observed in the WT.

[1] S. Tiwari, *et al.*, Photoacclimation strategies in cyanobacterial photosynthesis under dynamic light environments: implications in growth, fitness, and biotechnological applications. *Front Microbiol* 16:1686386. (2025). doi: 10.3389/fmicb.2025.1686386

[2] M.J. Mallén-Ponce, M. J. Huertas, A. M. Sánchez-Riego, F. J. Florencio, Depletion of *m*-type thioredoxin impairs photosynthesis, carbon fixation, and oxidative stress in cyanobacteria, *Plant Physiol* **187**, 1325–1340 (2021). https://doi.org/10.1093/plphys/kiab321
