## Supplementary material for "The cyanobacterium *Anabaena* uses pleomorphism as an acclimation strategy to high light stress": Table S1

| **Strain** | **Gene** | **Overexpression level**<br>OE- strain vs. WT<br>$\log_2$ fold change ± st dev |
|---|---|---|
| OE-*all2952* | *all2952* | 6.53±0.12 |
| OE-*all2981* | *all2981* | 3.49±0.63 |
| OE-*alr4579* | *alr4579* | 3.25±0.42 |
| OE-*alr5101* | *alr5101* | 8.26±1.75 |
| OE-*alr5324* | *alr5324* | 2.13±1.07 |
| OE-*alr5326* | *alr5326* | 9.72±1.07 |

**Table S1. Overexpression of genes encoding aPBPs.** The table indicates the aPBP-encoding genes that have been overexpressed in *Anabaena*, the name of the resulting strain and the overexpression level of the corresponding gene respect to the level in the WT.
