## Supplementary material for "The cyanobacterium *Anabaena* uses pleomorphism as an acclimation strategy to high light stress": Table S2

**Table S2.** Oligonucleotides used in this work

| **Name** | **Sequence** (5'-3') | **Comments** | **Experiment** |
|---|---|---|---|
| 3'-LeuS-F | GCTGGAATTCGTGTACCTGCCCACGATGTACG | For PCR amplification of 1645 bp from the 3' region of the *leuS* gene | Complementation of Δ*mre* mutants |
| 3'-LeuS-R | CATTGAATTAATCTCCTACTTAACCAACGACAAAATTAACTAAC | | |
| all0087-26 | TAAGTAGGAGATTAATTCAATGGGGCTTTTTAGGAACTTTCGC | For PCR amplification of the *mreB* ORF | Complementation of Δ*mreB* mutant<br>Probe for Northern assay |
| all0087-27 | ACGTGAATTCCTACATATTTCGAGATCGTCCGC | | |
| all0086-20 | TAAGTAGGAGATTAATTCAATGGTTACTGTACGTCGTTGGTGG | For PCR amplification of the *mreC* ORF | Complementation of Δ*mreC* mutant<br>Probe for Northern assay |
| all0086-21 | ACGTGAATTCCTAGTTGGACTTTTGTTGCTGTGAG | | |
| all0085-14 | TAAGTAGGAGATTAATTCAATGAAGACACCTTCATTTAACGG | For PCR amplification of the *mreD* ORF | Complementation of Δ*mreD* mutant<br>Probe for Northern assay |
| all0085-15 | ACGTGAATTCCTAACTTTCCAACATCTTCATTCTTTGC | | |
| all2952-F | AGGTTTACAAGCCGTGAACACC | | Probe for Northern assay |
| all2952-R | CATCGACTACAACGGGAGCATC | | |
| all2981-F | GGAGTAGGCCAGATAGCTGGC | | Probe for Northern assay |
| all2981- | CGCATACCACCCTTCAGCAG | | |
| alr4579-F | GGTCAACTGACGCAAGCAGTC | | Probe for Northern assay |
| alr4579-F | CCGCTTCACCACGCGATTG | | |
| alr5045-F | ATGACGGCGGGAATTAGTGC | | Probe for Northern assay |
| alr5045-R | GTACTAGCTGGAGGAAACGCAC | | |
| alr5101-F | AATGCGGCACACTAGTAAGGTG | | Probe for Northern assay |
| alr5101-R | CTGGTTGACGTAGGGCTTGC | | |
| alr5324-F | GCAGCCGCAAATCCGGACTTC | | Probe for Northern assay |
| alr5324-R | GCGTGGAGAGTACCAACCATTG | | |
| alr5326-F | TGTGACAGGAGGGTTAGTTGGC | | Probe for Northern assay |
| alr5326-R | CCCGTCAGGATAGCTGACTGG | | |
| RNA_5S_1F | TGGTACCACTCTGACCCCAT | | Probe for Northern assay |
| RNA_5S_1R | GGGCAACCCCTAGACTATCG | | |
| all2952_qPCR_1F | CCATAGAGGGAATGCCAATAG | | Q-PCR |
| all2952_qPCR_1R | GGTGTCAGAGAACGATTGAG | | |
| all2981_qPCR_1F | GTACTTCGTCGGGATTTGAG | | Q-PCR |
| all2981_qPCR_1R | CCGGAAACTGATCGCTAAA | | |
| alr4579_qPCR_1F | GTGGTTGCTTCTGAGGATAG | | Q-PCR |
| alr4579_qPCR_1R | GGGAGACTTGTTGGGTAATAG | | |
| alr5101_qPCR_1F | CAGCAGCCCGTACTTATTT | | Q-PCR |
| alr5101_qPCR_1R | CTTCAGTCGTTCCCAATGT | | |
| alr5324_qPCR_3F | CTGGGTAGGAAGAGACGATAA | | Q-PCR |

| | | | |
|---|---|---|---|
| alr5324_qPCR_3R | CTTGAAGTTCTCGACTGGTAAG | | |
| alr5326_qPCR_1F | CATCAGGTACTGTGAAAGAGG | | Q-PCR |
| alr5326_qPCR_1R | GCTCTAGGCGAATTGCTAATA | | |
| sfGFP-qPCR_2F | CTTGTCACTACTCTGACCTATG | | Q-PCR |
| sfGFP-qPCR_2R | TGTAGGTCCCGTCATCTT | | |
| all2952-2F | TTTCACACAGGAAACAGACCGTGGTAAGACAAAAAAAATTATTAAACCAG | | Fragment amplification for Gibson Assembly cloning |
| all2952-2R | TTGCTCATGGTGAATTCCATTTAGTTAGAAGAGGATGAATTAGGTCTAAG | | |
| all2981-2F | TTTCACACAGGAAACAGACCGTGTCGTCTAGGACTTTTGAAC | | Fragment amplification for Gibson Assembly cloning |
| all2981-2R | TTGCTCATGGTGAATTCCATTTAATTCGCCTTAGGACGCG | | |
| alr4579-2F | TTTCACACAGGAAACAGACCATGAATTCCCCCCAACCCCCTCA | | Fragment amplification for Gibson Assembly cloning |
| alr4579-2R | TTGCTCATGGTGAATTCCATTTATCTCGTAATTCTCCGCATATAATTT | | |
| alr5101-2F | TTTCACACAGGAAACAGACCATGCAATTCATATCTCTATCGCTTAC | | Fragment amplification for Gibson Assembly cloning |
| alr5101-2R | TTGCTCATGGTGAATTCCATTTAAGAATTACCAATCGAGAAACCCCT | | |
| alr5324-2F | TTTCACACAGGAAACAGACCGTGTCTTCATTAAGGACTTTTGAAGATAAG | | Fragment amplification for Gibson Assembly cloning |
| alr5324-2R | TTGCTCATGGTGAATTCCATTTATTTTTTATTCGGCCGAGGGC | | |
| alr5326-2F | TTTCACACAGGAAACAGACCGTGTCTTCATCAGAAAATATTGAACAGG | | Fragment amplification for Gibson Assembly cloning |
| alr5326-2R | TTGCTCATGGTGAATTCCATTTATTTCTTCTTTGGCTGAGGGC | | |
| all2952-1 | ATACTGCAGCACTCATGCCTACC | For PCR amplification of a region downstream of the fragment to be deleted | Construction of the Δ*all2952* deletion mutant |
| all2952-2 | AAGTTAGTTAGAAGATTGTCTTACCACCTG | | |
| all2952-3 | CAGGTGGTAAGACAATCTTCTAACTAACTT | For PCR amplification of a region upstream of the fragment to be deleted | |
| all2952-4 | ACTGCAGACCAAATCTATGGG | | |
| all2952-9 | ATCGCGCTTGCCTGGATCT | | Oligonucleotides used to check the structure of the *all2952* locus in the Δ*all2952* mutant |
| all2952-10 | TATTTTCACCAGACAAAAACCCGCAAG | | |
| all2952-11 | TGCAACTAAAACTGGTGATTTATCCCG | | |
